# Multi-Dimensional Structure and Dynamics Landscape of Proteins in Mammalian Cells Revealed by In-cell NMR

**DOI:** 10.1101/2022.09.20.508803

**Authors:** Harindranath Kadavath, Natalia Prymaczok, Cédric Eichmann, Roland Riek, Juan Gerez

## Abstract

Governing function, half life and subcellular localization, the 3D structure and dynamics of proteins are in nature constantly changing in a tightly regulated manner to fulfill the physiological and adaptive requirements of the cells. To find evidence for this hypothesis, we applied in-cell NMR to three folded model proteins and propose that the splitting of cross peaks are due to distinct structural states that arise from multiple target binding co-existing inside mammalian cells as a result of subcellular localisation including distinct cell compartments. In addition to peak splitting, we observed NMR signal intensity attenuations indicative of transient interactions with other molecules and dynamics on the microsecond to millisecond time scale.

---

The complexity of life is based on transient interactions between biomolecules within the complex environment of a cell. Many of the physical interactions of proteins with metabolites, nucleic acids, proteins and lipids have been confirmed^[1]^. However, investigating the structural aspects of such interactions in living cells, which are key towards a mechanistic understanding of biological processes, are highly challenging mainly due to the lack of a methodology suitable for such analyses. In-cell NMR spectroscopy, a method first reported in *E. coli* and then in eukaryotic cells^[2]^, has revealed interactions of various proteins with binding partners and ligands^[3]^, intracellular post-translational modifications, and the 3D structure of model proteins in the cellular environment. ^[2b,4]^

In order to investigate whether in-cell NMR can also provide insights into the different and multiple conformational states and dynamics of proteins in living mammalian cells, we produced in bacteria ^15^N-labeled protein G B1 domain (GB1), wild-type ubiquitin, and the second PDZ domain (PDZ2) of human tyrosine phosphatase hPTP1E, which were then purified and delivered by electroporation into mammalian cells^[2g,5]^, to a final concentration of ca. 5-10 µM delivering quasi background free NMR spectra (Fig. S20). 2D heteronuclear SOFAST-[^1^H,^15^N]-HMQC in-cell NMR experiments were performed to monitor for each residue its ^1^H-^15^N moiety yielding a dense net of probes in the protein. These spectra were then compared with the respective spectra of each protein in buffer. Compared to the spectra in buffer, the two most robust changes in the in-cell NMR spectra of all three systems are peak attenuations which were described in previous works and attributed to transient interactions with intracellular partners and dynamics on the microsecond (µs) to millisecond (ms) time scale ^[3a]^ and peak multiplication as discussed in detail below for each protein.

Our in-cell NMR experiments revealed that inside cells certain cross peaks of GB1 split into “daughter” peaks. We observe doubling, triplication and also higher multiplication states for many residues, termed further in this manuscript as ‘multiplicity’ of the peaks. While wide spread multiplicity is observed, all the multiple cross peaks are in close spectral proximity to the corresponding *in vitro* cross peak indicating the formation of the same fold but manifold perturbed by the cellular environment (Fig. 1a). Peak splitting is absent in the corresponding *in vitro* spectrum (Fig. 1a), where only one peak is observed for each residue, indicating that the intracellular milieu is responsible for the resonance multiplicity. For example, for Leu7 three cross peaks are observed in cells (Fig. 1a). The data is thus compatible with a model where in cells Leu7 of GB1 exists in three distinct structural states with an estimated life time between each other of longer than ms. Since the three peaks are similar in intensity and volume, these three structurally different species would be approximately equally present.

**Figure 1.**
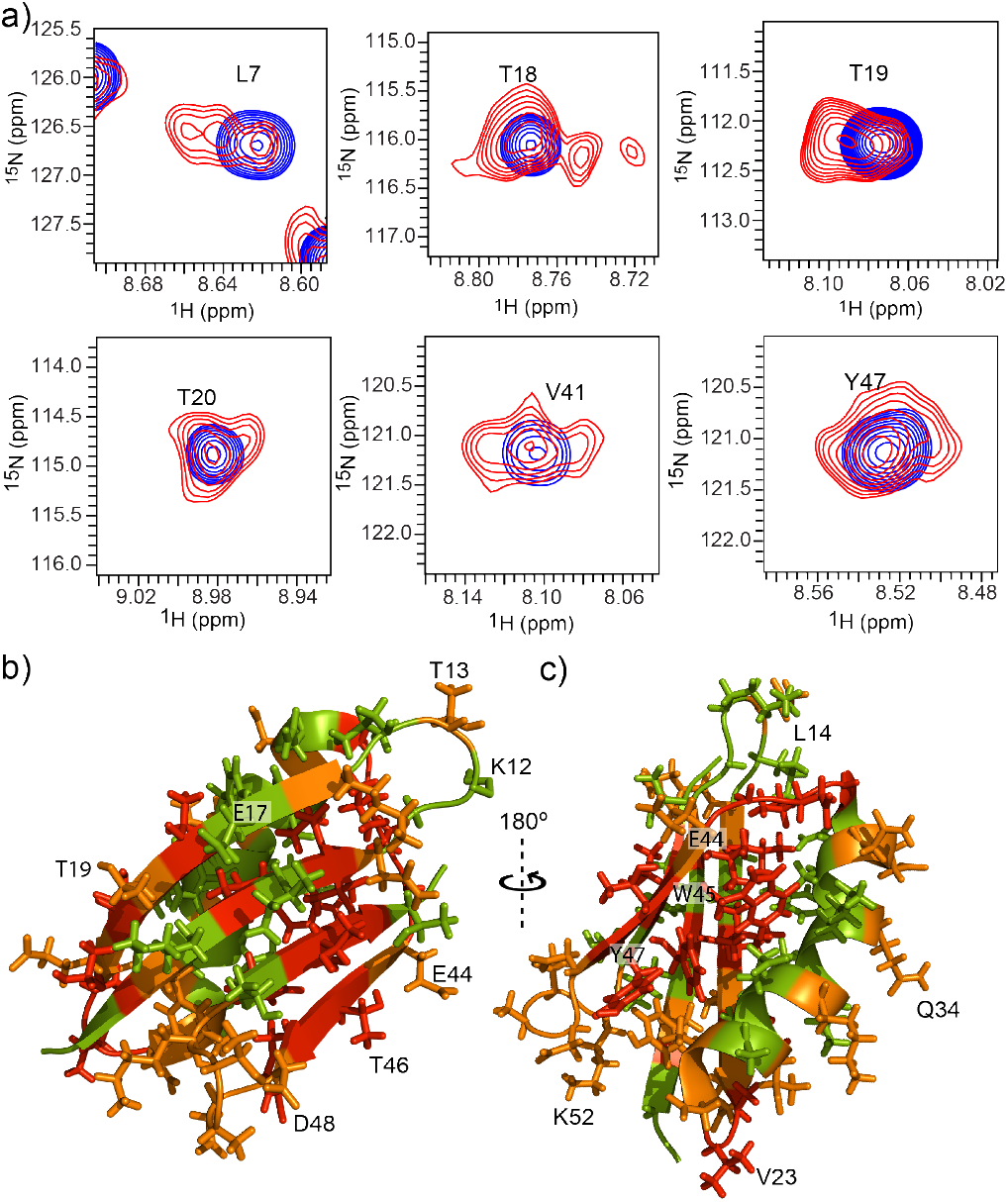
Peak splitting of GB1 in mammalian HEK-293 cells. a) Overlay of selected regions of the 2D SOFAST-[^1^H,^15^N]-HMQC spectra of ^15^N-labeled GB1 in mammalian cell HEK-293 (red) and in NMR buffer (blue) acquired at 310 K. The residues are labeled with the one letter code using the deposited data on BMRB 7280. b) and c) The multiplicities of GB1 resonances indicating the presence of distinct stable structural states are mapped onto the 3D structure of GB1 (PDB: 2N9L). Those resonances without any multiplicity are colored green, residues with multiplicity two are colored orange and those residues with multiplicity three or more are colored red.

Expanding this idea for Thr18 at least four long lived structural states are observed with some of them weakly populated (Fig. 1a). Supporting the notion of multiple structurally different states, peak multiplicity is observed for charged, polar, hydrophobic (including aromatics) side chains (Figs. 1b and 1c) as well as for both solvent exposed residues and in the hydrophobic core of the structure, in secondary structural elements and in loops (Figs. 1b and 1c). Peak multiplicity may result from specific and non-specific (transient) interactions with binding partners and endogenous components, but also from crowding resulting from a dense intracellular milieu or different environments of the subcompartmentalized cell interior (e.g. pH, ion composition). However, neither *in vitro* NMR experiments with the crowding agent Ficoll (Fig. S1), a pH titration from 7.5-5.5 (Fig. S2), nor salt titrations using NaCl and KCl from 0 to 150 mM (Figs. S3 and S4) yielded any peak multiplicity, wide spread chemical shift changes or peak intensity changes. Thus, peak multiplicity is not a mere consequence of different experimental conditions nor it is due to cell leakage and subsequent protein release during the 4 h NMR experiment (Fig. S21a and S21b) or due to protein degradation (Fig. S21c) but rather results from interactions with intracellular molecules. To support this notion, we acquired spectra of GB1 in the presence of cell lysate of HEK-293 cells, with a protein concentration of ∼5 µM, and compared them with the corresponding spectrum acquired in living cells (Figs. S5, S6 and S7). The cell lysate was separated into the soluble fraction and the pellet which contains membranous organelles such as nuclei, mitochondria and plasma membrane components. Again we observed residue-specific multiplicities both in the soluble fraction and in the pellet. Interestingly, the new peaks add up only partially with the in-cell NMR cross peaks. For example, residues Lys30, Gly40, Gly43, and Ala50 show multiplicity in cells, but remained unchanged both upon addition of soluble fraction or pellet to GB1, suggesting that they are only sensitive to an intact cell. However, residues Leu9 and Ala26 sum up to the in-cell spectra suggesting that these residues interact with either soluble molecules as well as molecules contained in organelles (Figs. S6 and S7).

In the next attempt we tested whether the origin of the peak multiplicity is due to distinct subcellular localization by delivering into the cells a GB1 variant containing a mitochondria-anchoring tag and studied its behavior both at 1 and 4 h post electroporation by confocal microscopy and NMR. If confirmed the peak multiplicity hypothesis, GB1 association to mitochondria would result in changes in peak multiplicity due to the absence of other forms of GB1 present in the homogenously distributed wild-type protein. We found that at 1 h GB1 was widely distributed inside the cells, indicating that the protein had been efficiently delivered but not yet reached its final destination. In contrast at 4 h this protein was mostly found in mitochondria (Fig. S8). Interestingly, the peak multiplicity present at 1 h is largely reduced at 4 hours post electroporation (Fig. S9). The GB1 association to mitochondria yielded the reduction of peak multiplicity (Fig. S9) attributed of a predominant mitochondria-specific structural species of GB1 in these cells as well as the existence of several other structural states of GB1 in other compartments.

To get insights into the origin and specificity of the interactions yielding peak multiplicity two additional experiments were undertaken. First, *in vitro* it is demonstrated that the NMR spectrum of GB1 as well as ubiquitin (studied below) is largely insensitive to metabolites and other small molecules purified from a cell extract by filtration (Figs. S10 and S11) requesting rather proteins, RNA, or/and membrane entities to be binding ligands. Second, peak multiplicity in cells is followed by a concentration series of GB1 (4, 10 and 40 µM) (Figs. S12 and S13). The loss of peak multiplicity with concentration towards a single resonance close to the *in vitro* resonance indicates the saturation of the binding partners with estimated µM binding affinities.

In addition to peak multiplicity, also peak attenuations of the GB1 resonances are observed (Fig. S14) indicating the presence of transient interactions with other molecules on the µs to ms time scale. In particular the cross peaks of the loops (e.g. Lys12-Leu14 and Asp48-Lys52) and of solvent exposed residues (e.g. Glu17, Val23, Glu44) are highly affected by the cellular environment while the hydrophobic core (with the exception of the partially solvent exposed Trp45, Tyr35 and Tyr47) seems not to be altered. Overall, these studies indicate that GB1 interacts in cells in part transiently with other biomolecules in the µM affinity range organelle-distinct.

We next analyzed human wild-type ubiquitin, a mammalian protein with multiple confirmed specific interactions in the cell^[6]^, in Cos7 cells. Even more pronounced peak multiplicity was found on ubiquitin (Fig. 2a). Lys6 and Ser65 show around 10 cross peaks in the in-cell NMR spectrum. As with GB1, the multiplicity of residues includes charged (e.g. Lys6, Glu8), polar (e.g. Thr9, Gln49) as well as hydrophobic residues (e.g. Val26, Leu43, Lleu50). When mapped onto the 3D structure of ubiquitin (Figs. 2b and 2c) multiplicity is observed for both solvent exposed and hydrophobic core residues (e.g. Leu15, Val17, Val26, Ile30, Leu67), β-strands (e.g. Lys11-Glu18) as well as helical secondary structures (e.g. Val26-Gly35). The highest multiplicity is observed on the C-terminal half of the α-helix as well as for β-strand β2 (comprising residues Thr9-Val17) and spatially adjacent residues. In addition to being covalently linked via its N-terminus to other proteins in a process called ubiquitination, ubiquitin can also bind non-covalently to other proteins such as the proteasome, ZnF, PLIC1, UBCH5C proteins and E3 ubiquitin ligases (Fig. S15). Previous in-cell NMR studies have shown evidence for intracellular interactions of ubiquitin.^[2a,2b,7]^ Most of the known non-covalent interactions are through the β-sheet in the vicinity of Ile 44 and do not superimpose the multiplicity observed (neither with β-strand β2 nor with the α-helix) with the exception of the binding with the VHS domain which shows a secondary binding site around K11, I13, K27, K29, K33 and K63^[8]^. Noteworthy is the high multiplicity of all the ubiquitination-relevant Lys residues (i.e. K6, K11, K27, K29, K33 and K48) but K63 is important in polyubiquitination^[9]^. This may hint to the origin of the observed structural plurality of ubiquitin in cells to be ubiquitination. In support of this hypothesis is the close qualitative resemblance of the chemical shift changes on ubiquitin upon ubiquitination documented by McKenna et al.^[10]^ and the ones documented here.

**Figure 2.**
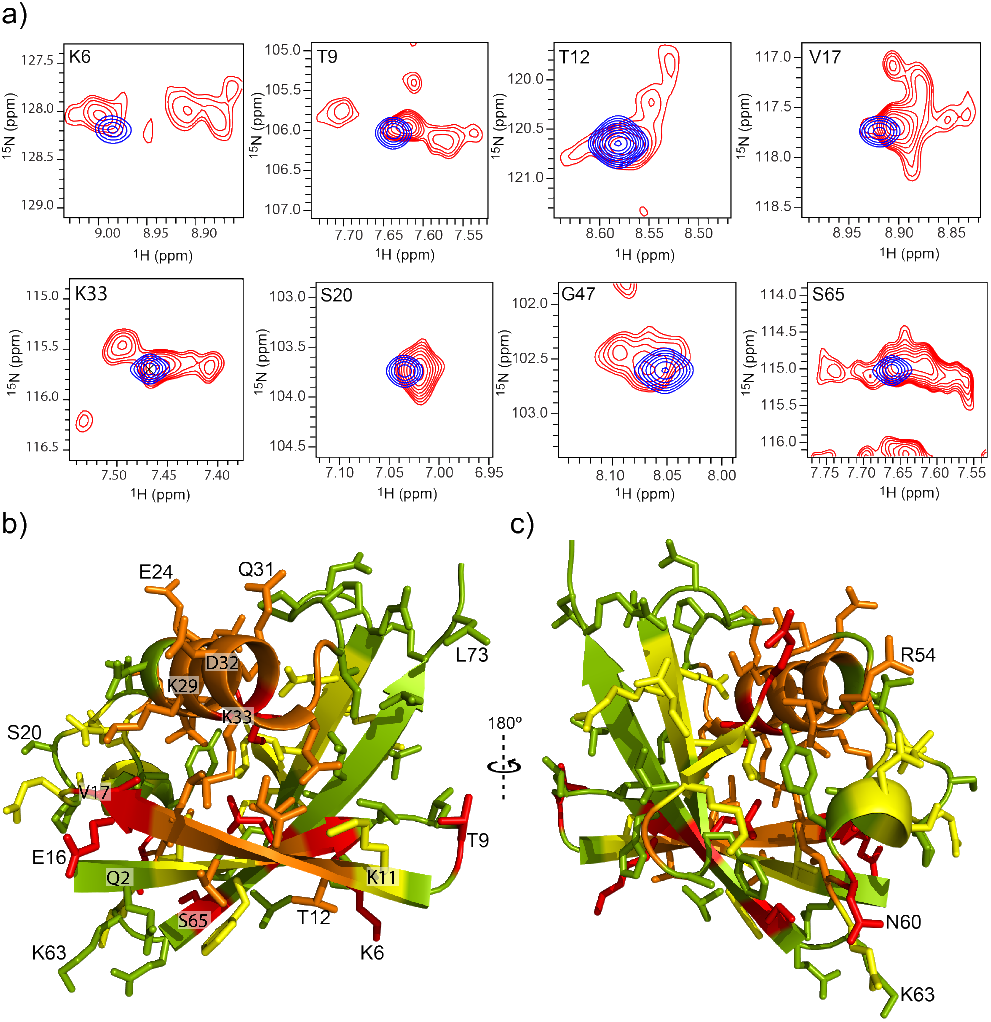
Peak splitting of ubiquitin in mammalian Cos7 cells. a) Overlay of selected regions of the 2D SOFAST-[^1^H,^15^N]-HMQC spectra of ^15^N-labeled ubiquitin in Cos7 cells (red) and in NMR buffer (blue). The residues are labeled with the one letter code using the deposited data on BMRB. b) and c) The multiplicities of ubiquitin resonances indicating the presence of distinct stable structural states are mapped onto the 3D structure of ubiquitin (PDB: 1UBQ). Those resonances without any multiplicity are colored green, residues with multiplicity two are colored yellow, with multiplicity three, orange and those residues with multiplicity four or more are colored red, respectively.

In order to strengthen the indicated origin of the peak multiplicity to specific interactions of ubiquitin with its binding partners we studied by in-cell NMR an inactive ubiquitin lacking the C-terminal two Gly residues which are essential for ubiquitin attachment to target proteins.^[11]^ This protein variant was delivered into mammalian cells and compared with its wild-type counterpart (Fig. S16). Indeed, an almost complete loss of the peak multiplicity is observed. Furthermore, the cross peaks superimpose with the *in vitro* reference cross peaks indicating the inert nature of the inactive mutant in cells.

As in the case of GB1, in addition to the peak multiplicity, signal loss in the in-cell NMR spectrum of wild-type ubiquitin when compared with the *in vitro* spectrum is observed indicating again also the presence of transient interactions (Fig. S17). Note that all the five β strands of ubiquitin which are alternatively anti-parallel and constitute more than half of the 3D structure are significantly affected. In particular, the loops 1, 2 and 4 with residue stretches Thr7-Gly10, Val17-Ile23, Phe45-Lys48 undergoes intermediate exchange as evidenced by the reduction in their volume ratios (Fig. S17). Surprisingly the residues showing transient interactions or/and dynamics induced by the cellular milieu have mainly solvent exposed side chains. In addition to the loops 1, 2 and 4 certain residues in loop 6 such as Lys63 and Glu64 are highly affected by the cellular environment suggesting the involvement of Lys63 in the ubiquitination and transient interactions.

We next analyzed the second PDZ domain (PDZ2) from human tyrosine phosphatase hPTP1E^[12]^ in A2780 cells. PDZ domains mediate the interactions of several proteins along the signal transduction pathways.^[13]^ We again observed multiplicity of PDZ2 resonances with up to ten cross peaks, for instance for His71 and Lys 72 (Fig. 3a, Fig. S18). Of note PDZ2 displays high multiplicity on a particular set of amino acid residues around the canonical protein-ligand pocket i.e. Gly19, Ile20, Ser21, Val22, Gly23, Gly24 of the β-strand β2 and residues Thr70, His71, Lys72, Ala74, Val75, Leu78, Arg79 and Thr81 of the α-helix (Fig. 3c, d). In addition, the short helix including Glu47-Gly50 as well as the β-strand β4 spanning from Arg57-Ala60 both far in space from the ligand binding site show for each residue multiplicity. With time, cross peaks of residues mostly around His71 change and is attributed to a metabolisis-dependent pH drop in the sample (Fig. S22).

**Figure 3.**
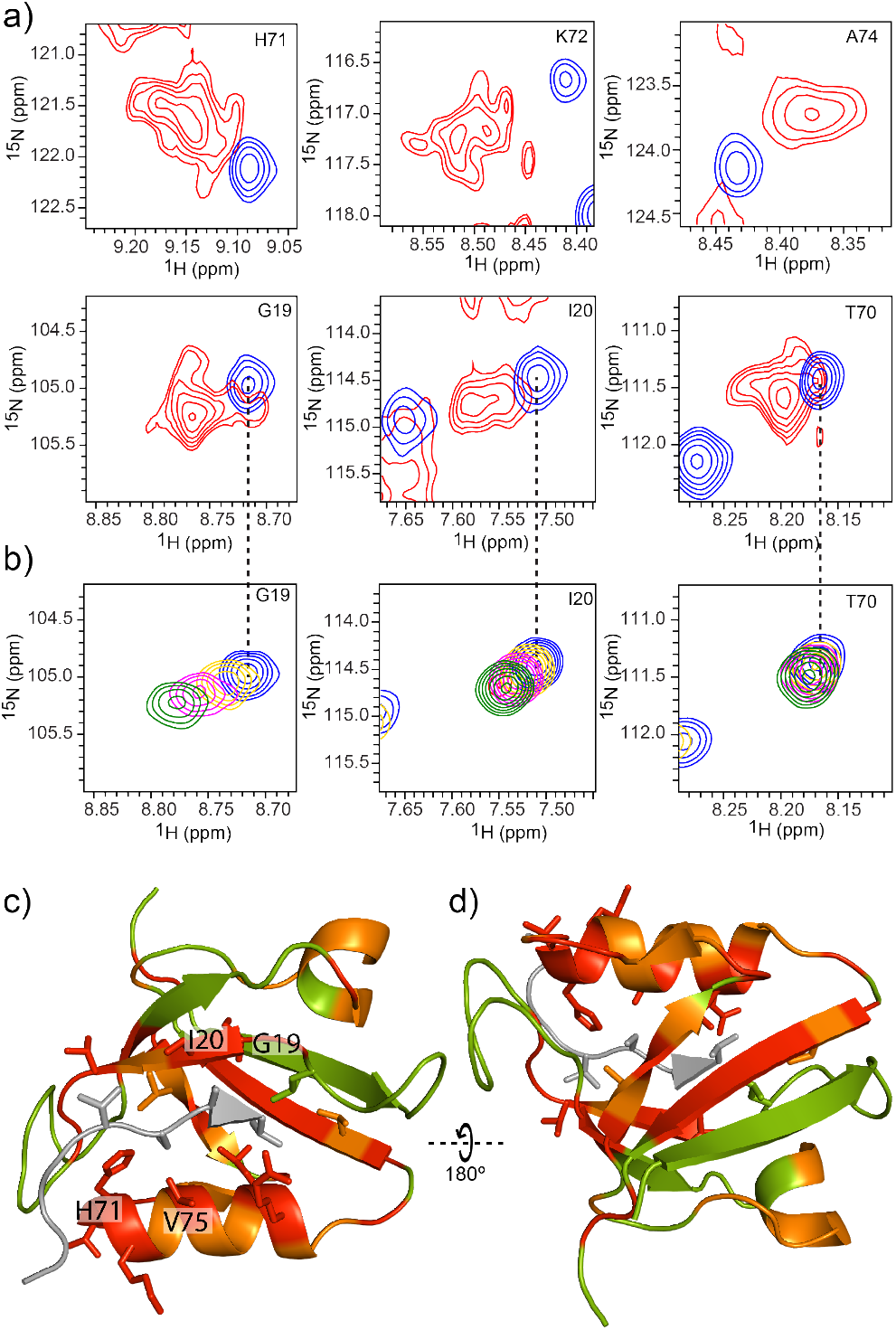
Peak splitting of the PDZ2 domain in mammalian A2780 cells. a) Overlay of selected regions of the 2D SOFAST-[^1^H,^15^N]-HMQC spectra of ^15^N-labeled PDZ2 in A2780 cells (red) and in NMR buffer (blue). b) Ligand-binding NMR titration of the ^15^N-labeled PDZ domain with unlabeled peptide VSAV. Overlay of selected regions of the 2D SOFAST-[^1^H,^15^N]-HMQC spectra of ^15^N-labeled PDZ are shown in absence of ligand (blue cross peaks), in a 1:1 (yellow), 1:2 (magenta) and 1:3 (in green) PDZ : ligand ratios. c, d) 3D structure of the PDZ2 domain in complex with a peptide ligand (PDB: 1D5G) with the residues color coded according to the multiplicity of peaks described in the caption to figure 1 and the ligand peptide colored in grey. Selected residues are labeled.

It is straight forward to assume that peak multiplicity represents the multi structural states caused by ligand binding. In full support of this, an *in vitro* NMR titration of the peptide VSAV from the C-terminus of guanine nucleotide exchange factor being a known binding partner of the PDZ2 domain under study shows similar chemical shift perturbations as the in-cell spectrum for Gly19, Ile20, and Thr70 (Fig. 3b). However due to the presence of a single peptide entity, only one cross peak is observed. It is expected that in cells PDZ2 binds to many different protein segments present in the cell yielding distinct structural states detected by in-cell NMR. Since the *in vitro* studies show binding of the peptide in the fast exchange regime (i.e. µs to ms time scale) with a K_D_ in the µM range, the distinct structural states shown by in-cell NMR are likely also due to transient interactions, but each interaction spatially separated from the others and thus of local nature. This conjecture is supported by the finding that peak intensity attenuation in cells when compared with the *in vitro* spectrum indicative of transient interaction coincides with the structural area for which peak multiplicity is observed (Fig. S19). Furthermore, *in vitro* analysis demonstrated the presence of protein allostery in the PDZ2 domain between the ligand binding pocket and the short helix comprising Glu47-Gly50 and β-strand β2 comprising residues R57-A60.^[14]^ That the same segments show multiplicity in the in-cell NMR spectra indicates that protein allostery between the binding pocket and these two remote sides is not only present in cells but is also influenced by ligand interaction.

In summary, the in-cell NMR data presented here reveals the presence of multiple either long-lived or spatially-separated short-lived structural and dynamical states for the three proteins studied when present inside mammalian cells. These multi-dimensional structure/dynamics landscape is thereby directed by the cellular environment likely containing both specific interactions, as demonstrated in the case of the ubiquitin and PDZ2 domain on ligand-protein binding, as well as unspecific interactions (namely quinary interactions)^[15]^ as observed for the non-mammalian protein GB1. The large plurality of distinct structures around the same fold is thereby expected to be key for the achievement of the required complexity of a functional cell as highlighted by the potential finding of a ligand-induced allosteric effect on the PDZ2 domain when present in the cell. The structural biology inside a cell is just emerging^[16]^. It will likely compile a large richness in information necessary if a comprehensive understanding on the cellular life is aimed at.

