## Supporting Information for "Multi-Dimensional Structure and Dynamics Landscape of Proteins in Mammalian Cells Revealed by In-cell NMR"

### Experimental Section

*Sample preparation*

All proteins used here were recombinantly expressed in *E. coli* BL21(DE3) cells in M9 minimal media, supplemented with $^{15}NH_4Cl$, D-($^{13}$C)-glucose (if needed) and appropriate antibiotics and induced overexpression with isopropyl-b-D-thiogalactoside (IPTG). We used [U-$^{15}$N] labeled GB1, wild-type human ubiquitin (accession number 1YX5_B) and PDZ2 domain. [U-$^{15}$N,$^{13}$C] labeled PDZ2 domain was used for the resonance assignment.

*Cell culture*

HEK-293, Cos7 and A2780 cells were purchased and cultured in Dulbecco's Modified Eagle Medium (DMEM, pH 7.4) supplemented with 10% fetal calf serum, 4 mM L-glutamine, 100 U/ml penicillin and 100 mg/ml streptomycin.

*In-cell NMR*

Up to 10 plates of 90% confluent HEK-293, Cos7 and A2780 cells were used for one in-cell NMR experiment. For in-cell NMR experiments the cells were grown in DMEM, pH 7.4 supplemented with 10% fetal calf serum, 4 mM L-glutamine, 100 U/ml penicillin and 100 mg/ml streptomycin for 2 days, then harvested, pooled, centrifuged at 200 x g for 3 min, and kept at 37°C for 10 min as cell pellets. An aliquot of 5-20 mg of protein was re-suspended in 100 µl of sterile PBS pH 7.4 (GIBCO). The protein sample in PBS was electroporated (NEON transfection system) or mixed at a 1:1 ratio with electroporation buffer (120 mM phosphate, 5 mM KCl, 15 $MgCl_2$, 50 mM NaCl, pH 7.5) (AMAXA-LONZA). Cells were kept for 5 min in the electroporation mixture containing labeled protein and finally subjected to electroporation using a NEON (Thermo Fischer) or Nucleofector-IIb (AMAXA-LONZA) electroporator. Immediately after electroporation, cells were washed twice with pre-warmed media, plated, and incubated at 37 °C for 4 hours. Mitochondrial GB1 was analysed also at 0 h post electroporation. After this recovery period, the cells were harvested by mild trypsinization, washed once with pre-warmed media, and once with PBS pH 7.4 containing 5% D2O.

*NMR Spectroscopy*

NMR experiments were performed on a Bruker Avance NEO 700 MHz and Avance III HD 600 MHz spectrometers equipped with a cryogenically cooled proton-optimized $^{1}$H[$^{13}$C/$^{15}$N] TCI probe. Bruker Topspin 2.1, 3.2 and 4.0 were used for data acquisition. Specifically, 2D $^{15}$N-$^{1}$H SOFAST HMQC spectra were acquired with a data size of 128 × 512 complex points for a sweep width (SW) of 28.0 ppm ($^{15}$N) and 16.7 ppm ($^{1}$H), 512 scans, 60 ms recycling delay (acquisition time ~ 4 h). All the *in vitro* NMR experiments were performed in NMR buffer (PBS, pH 7.4) and at temperatures of 283K, 298 K and 310 K.NMR spectra were processed with either Bruker Topspin 4.0 and NMRFAM-Sparky[1] or PROSA[2] and CARA.[3] Visualization and data analysis were carried out in NMRFAM-Sparky[1] or CARA.

GB1 and ubiquitin NMR spectra were analyzed with the backbone resonance assignment published previously (BMRB: 7280 and 6457, respectively). The backbone resonance assignment of PDZ2 domain was performed in house using 3D HNCA, HNCOCA, HNCACB, and CBCACONH spectra.

*Titrations & quantifications*

The salt titration experiments of GB1 were performed in the presence and absence of NaCl and KCl, where the ionic strength was 0 and 150 mM. PDZ2-ligand titration was carried out using 100 uM PDZ2 domain containing 0, 100, 200 and 300 $\mu$M VSAV peptide ligand. A 36 mM ligand stock in NMR buffer was used. To perform experiments with cells lysates, HEK-293 cells were lysed and the centrifuged at 10.000 xg for 30 min. The supernatant containing soluble proteins and the pellet were separated and used for NMR determinations as soluble and pellet fractions. To these samples ~5 $\mu$M $^{15}$N-labeled GB1 was added and acquired NMR spectra.

Quantification of in cell protein levels were determined by using a 1D [$^{15}$N,$^{1}$H] HMQC NMR experiment of the given protein in vitro with known concentration and compared it with the corresponding transexpressed protein signal in mammalian cells (with focus on the strongest signals) possible because NMR is a quantitative method.

*Generation of mitochondrial GB1 and delta-di-glycine ubiquitin*

The bacterial expression plasmid encoding mitochondrial GB1 was created by PCR. At the N-terminus of mitochondrial GB1 we included an 6XHis tag for niquel purification and a TEV site to remove the 6XHis tag from the purified protein. At the C-termius GB1 contained the FLAG tag for immunostaining, and the peptide LSLRQSIRFFKPATRTLCSSRK (to deliver GB1 tomitochondria). The plasmid encoding wild type GB1 was used as template of PCR and the primers were:

Forward:AGC**CATATG**CTGAGCCTGAGGCAGAGCATCAGGTTCTTCAAGCCCGCCACCAGGACCCTGTGCAGCAG CAGGCAGTACAAGCTTATCCTGAACGGTAAAACC,

Reverse:TAT**GGATCC**CTGCTGCTGCACAGGGTCCTGGTGGCGGGCTTGAAGAACCTGATGCTCTGCCTCAGGCTC AGCTTGTCGTCGTCGTCCTTGTAGTC

The PCR product was then purified and digested with NdeI and BamHI (underlined in the primers), and finally ligated into a pET11a vector digested with the same endonucleases.

The bacterial expression plasmid encoding delta-di-glycine ubiquitin was created by PCR. At the N-terminus of delta-di-glycine ubiquitin we included a FLAG tag for immunostaining. The plasmid encoding wild type ubiquitin was used as template of PCR and the primers were:

Forward: AGCCATATGGACTACAAGGACGACGACGACAAGCAGATCTTCGTCAAGACG,

Reverse: GTAGAATTCTCAACGTAGACGTAAGACAAG

The PCR product was then purified and digested with NdeI and EcoRI (underlined in the primers), and finally ligated into a pGEX vector digested with the same endonucleases.

*Confocal microscopy*

Cells were grown in glass coverslips for the indicated times and then fixed; the cells were rinsed 3 times in Mg-PBS (PBS supplemented with 10 mM $MgCl_2$), treated with blocking solution (5% BSA fraction five in Mg-PBS) and then incubated for 10 min in Mg-PBS plus 1% Tritton X100. After 3 rinses in Mg-PBS the coverslips were incubated in the dark for 4 hours with the antibodies. Then the coverslips were washed three times in Mg-PBS, and the in fluorochrome-conjugated secondary antibodies were added for 1 h. After that, the coverslips were rinsed with Mg-PBS containing 1 % DAPI, washed again and coverslipped with mounting medium (Immu-Mount, Thermo scientific). Fluorescent staining was visualized with a confocal Zeiss Spinning Disk confocal microscope.

*Metabolite enrichment*

10 plates of 90% confluent HEK-293 cells were grown in DMEM, pH 7.4 supplemented with 10% fetal calf serum, 4 mM L-glutamine, 100 U/ml penicillin and 100 mg/ml streptomycin for 2 days. The cells were washed twice with PBS, then harvested, pooled, centrifuged at 200 x g for 3 min, and kept at -80°C for 10 min as cell pellets. Four cycles of frezze/thawing and three cycles of sonication were used to lyse the cells. The lysed cells were centrifuged at 4 C for 30 min at 10.000 x g and the supernatant was filtered using a 2 kDa cut off filter.

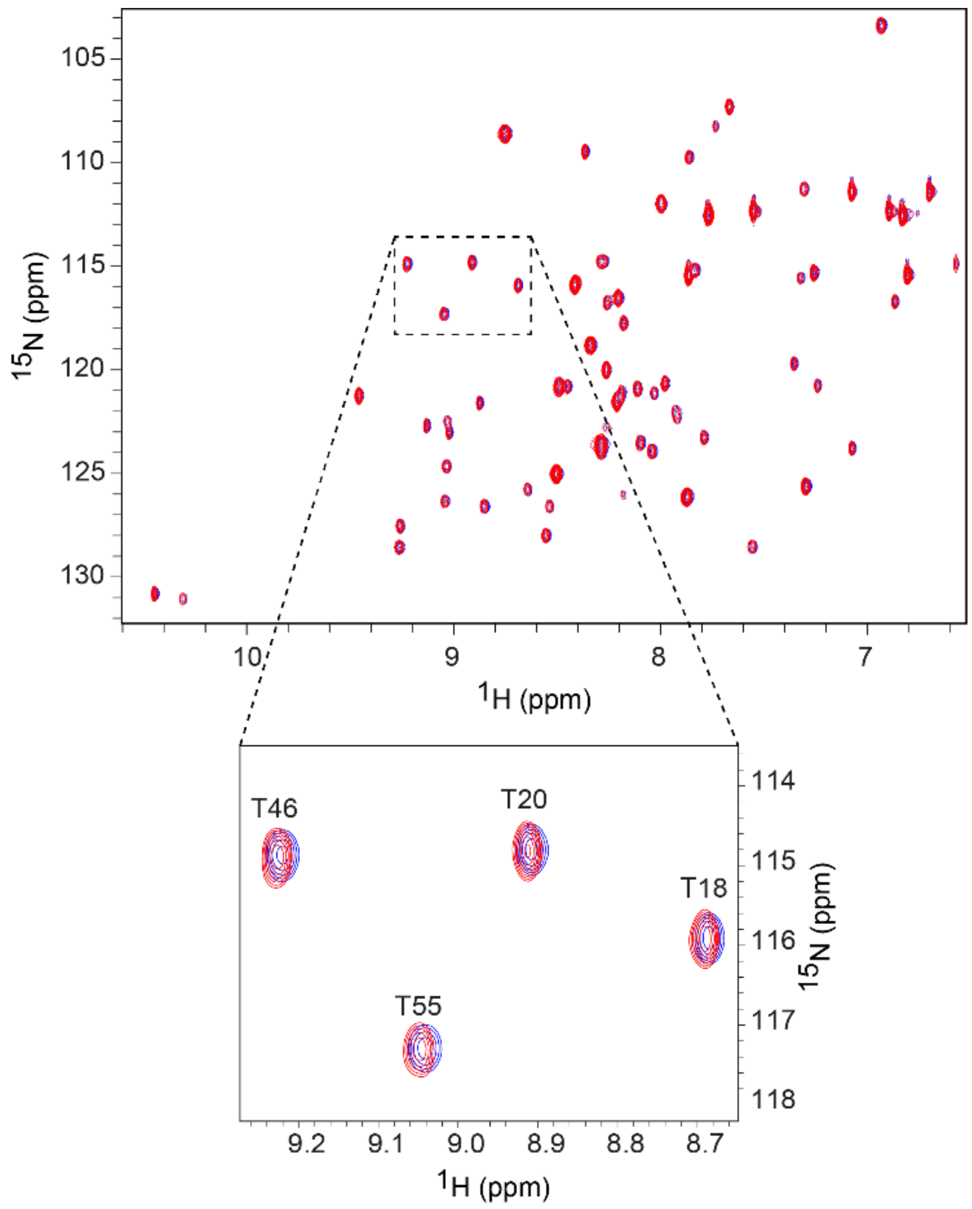


**Fig S1. Influence of crowding on the GB1 resonances.** Comparison of the spectra acquired in buffer (blue) and in presence of 200 mg/ml ficoll (red). Notice that the crowding agent ficoll does not induce any detectable peak splitting on GB1.

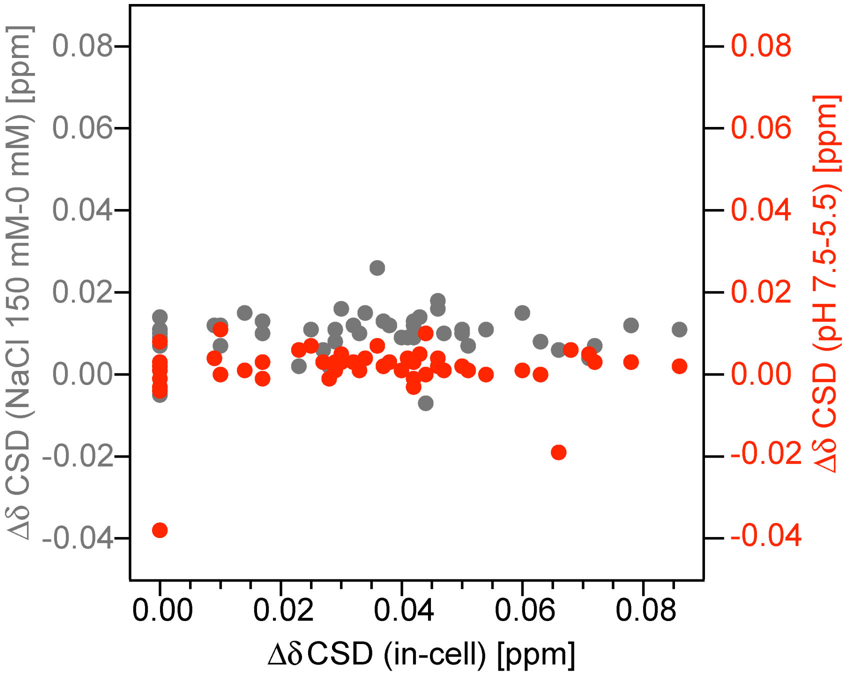


**Fig S2. Effect of pH and salt concentration on the GB1 resonances**. The extent of chemical shift dispersion of each residue (taken in x-axis) is correlated with the chemical shift perturbation induced as a result of salt titration using NaCl (left y-axis) and chemical shift perturbation due to pH titration (right y-axis).

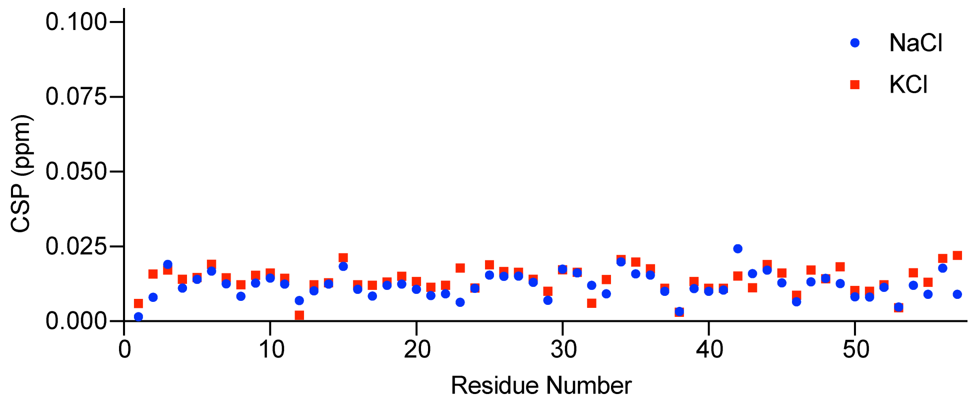


**Fig S3. Chemical shift perturbation (CSP) of GB1 resonances induced by change in ionic strength.** The CSP of GB1 resonances as a result of 150 mM NaCl (blue) and 150 mM KCl (red) with respect to 0 mM respective salt. The given CSP is the combined $^{1}H$ and $^{15}N$ shifts.

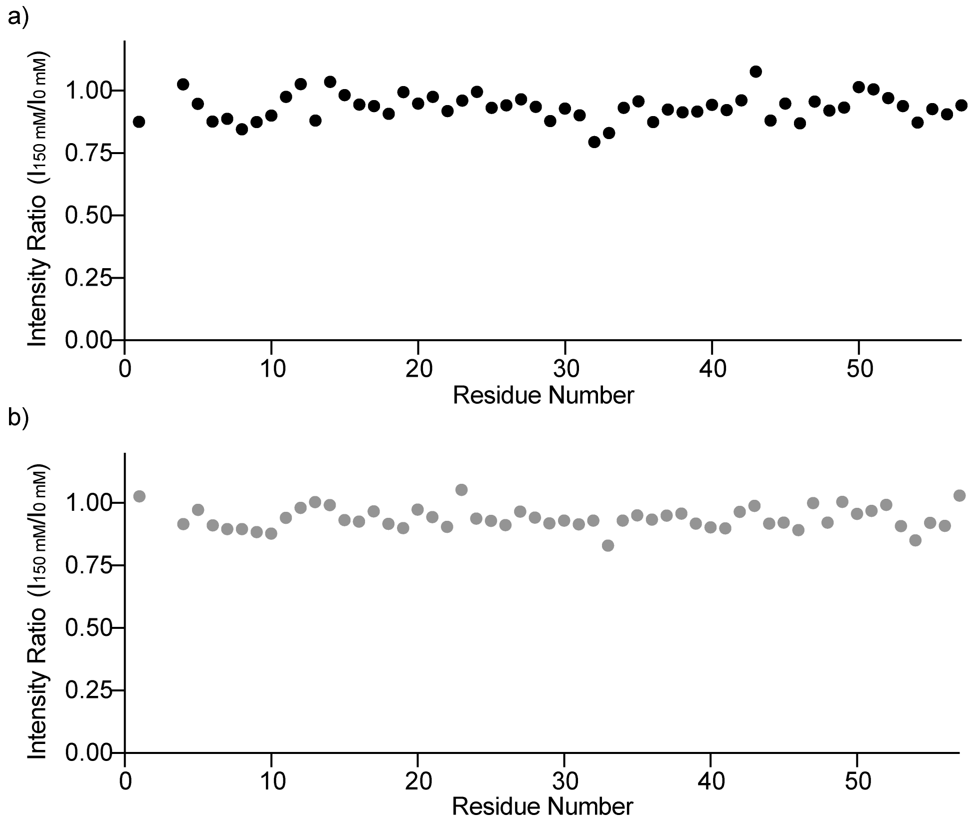


**Fig S4. Effect of salt concentration on the intensity of GB1 resonances**. The variation in intensity of GB1 resonances as a result of high salt concentration using NaCl (a, black) and KCl (b, grey). Intensity ratio was calculated by dividing the intensity of GB1 peaks at 150 mM NaCl or KCl concentration by the intensity of GB1 peaks at 0 mM NaCl or KCl.

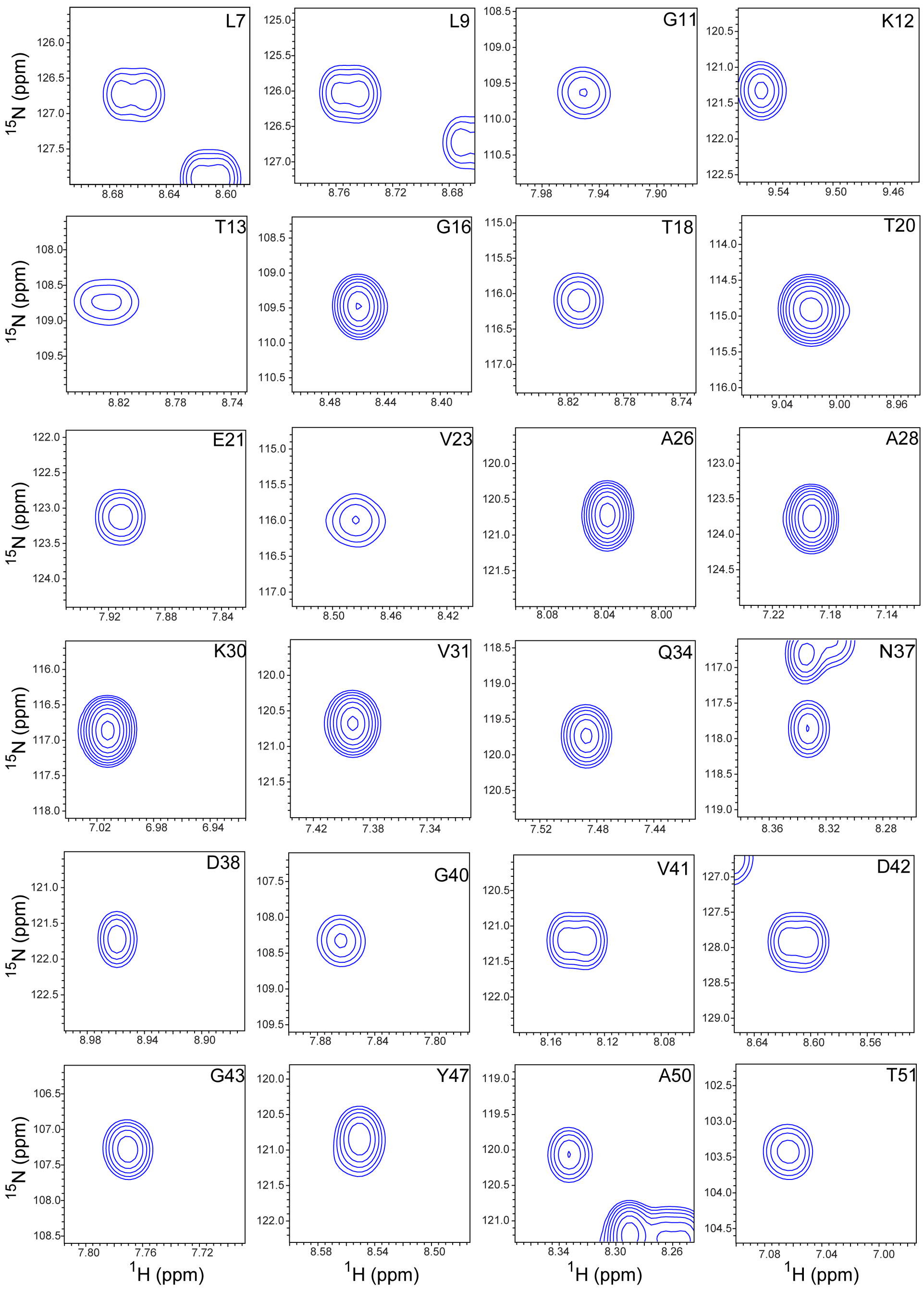


**Fig S5. Single peak for each resonance for GB1 in vitro.** Selected GB1 NMR spectral resonances acquired in NMR buffer (PBS, pH 7.4). The resonance assignment of the peak is shown in the upper right corner of the spectral region with one letter amino acid labels.

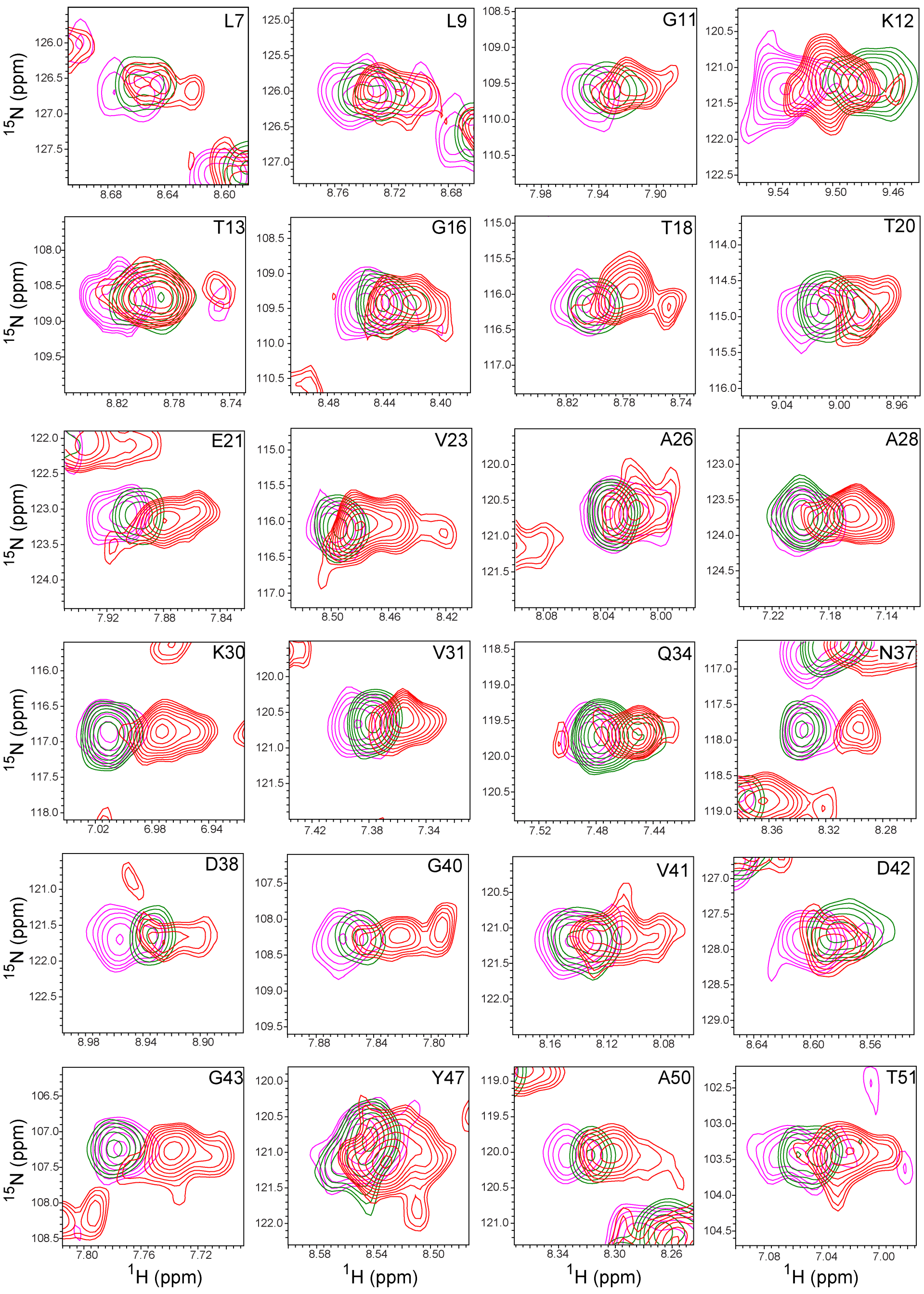


**Fig S6. The multiplicity of the GB1 cross peaks in cells can only be reconstituted in part by ex situ measurement with GB1 added to cell lysate or cell pellet**. Comparison of GB1 cross peaks acquired in living HEK-293 cells (red), in the soluble fraction of the HEK-293 cell lysate (green) and in presence of cell pellet (magenta). The shown selected resonances are in accordance with Fig S5. For many residues the cross peaks of the spectra recorded in cell's do not add up of the cross peaks from the cell pellet and soluble fraction of the cell lysate indicating the in cell NMR spectrum being sensitive to the intact nature of the cell, while for some residues a partial sum up is observed (for example A26, L9).

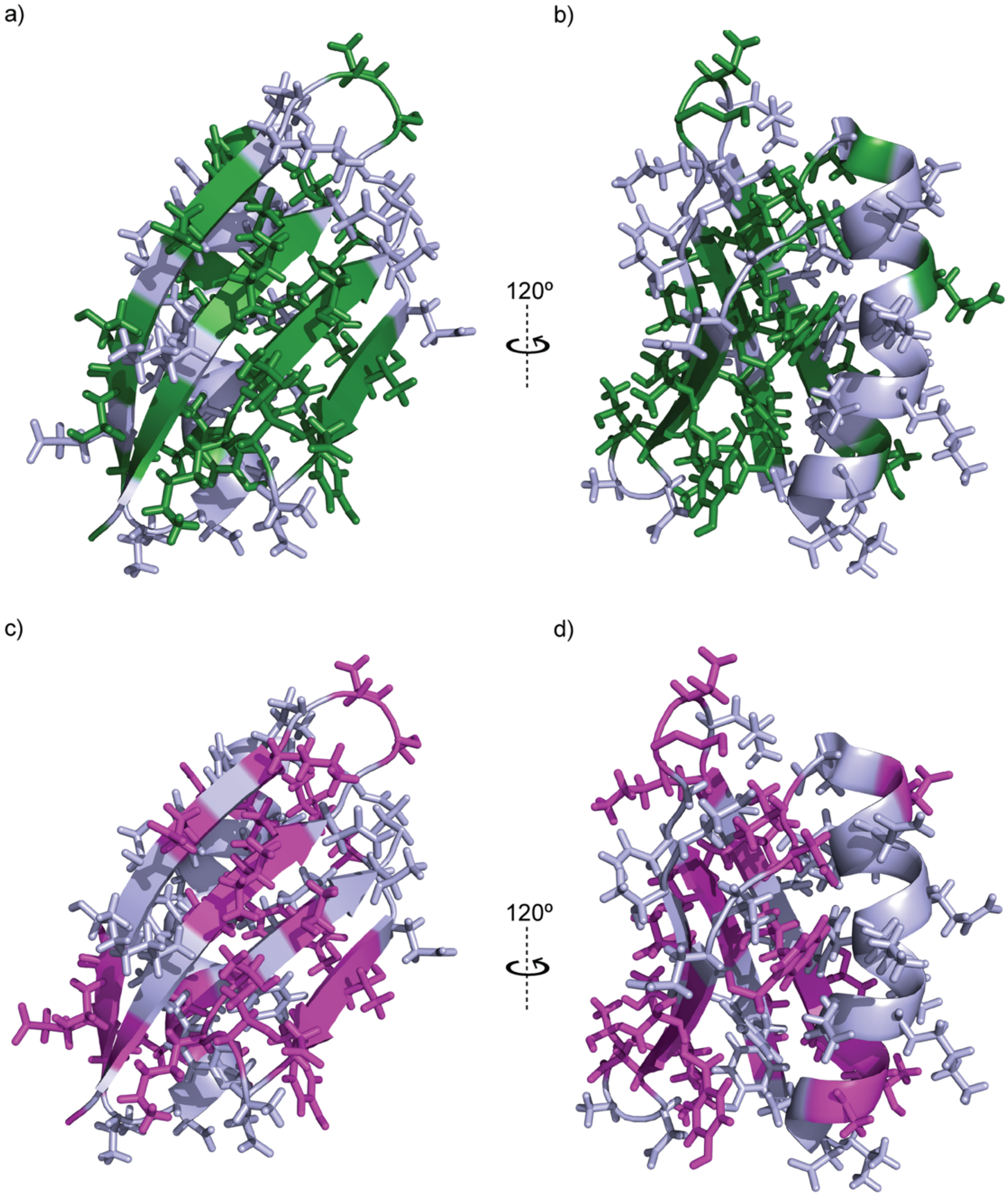


**Fig S7. Multiplicity of cross peaks upon addition of GB1 into (a,b) cell lysate and (c,d) into cell pellet mapped onto the 3D structure**. a) and b) Residues showing multiplicities of GB1 resonances indicating the presence of distinct stable structural states in HEK-293 cell lysate are mapped onto the 3D structure of GB1 (PDB: 2N9L). Those residues with multiplicity two or more are colored green and those residues without any multiplicity are colored light blue. c) and d) Same structure and orientation as in a and b respectively. Multiplicities of GB1 resonances in the presence of cell pellet are colored purple and those residues without any multiplicity are colored light blue.

a)

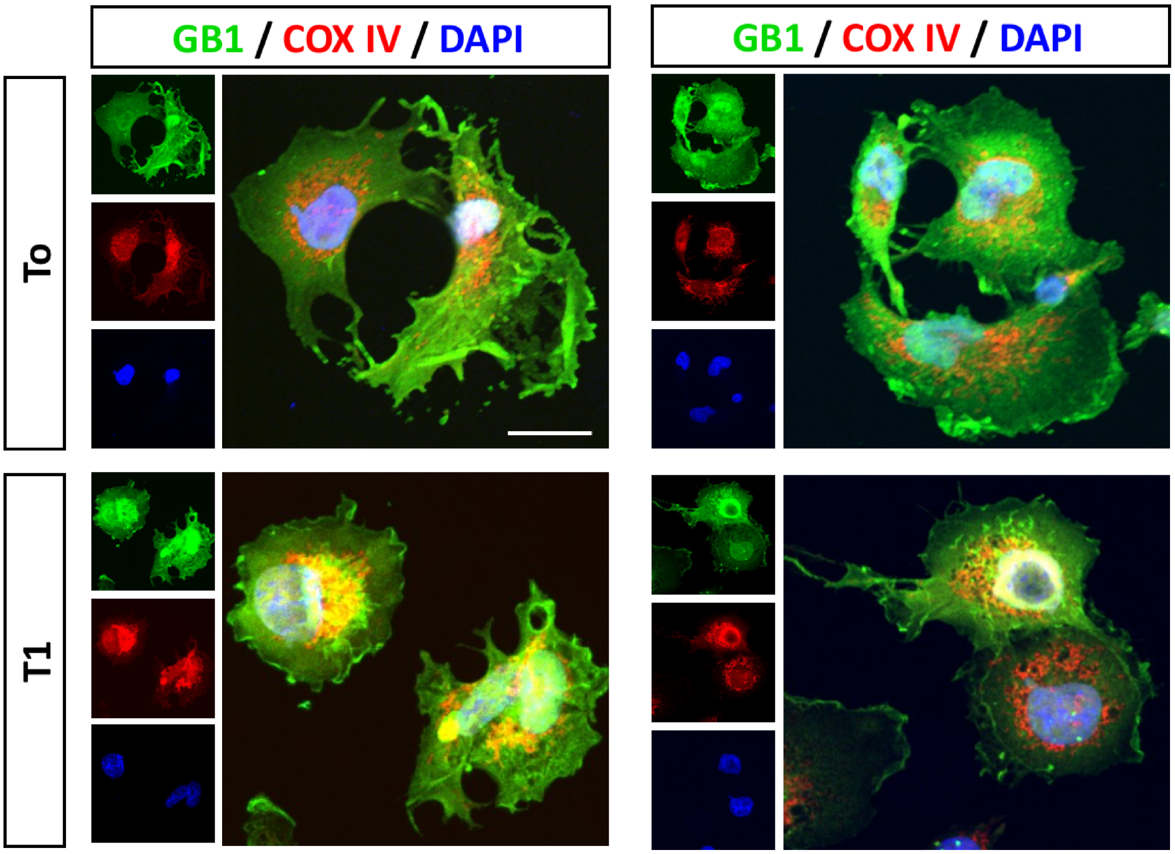


**Fig S8. Localization of mitochondrial-tagged GB1.** Representative confocal microscopy images of GB1 (green) containing a mitochondrial tag electroporated into Cos7 cells. (a) The image taken 1 h after electroporation (T0), indicates that GB1 is found throughout the cell. (b) The image taken after 4 hours (T1) revelas the re-localization of GB1 to mitochondria (red). For this experiment we used Cos7 cells that, compared to HEK-293 cells, are bigger in size allowing to visualize the re-localization to mitochondria more precissely. Immunofluorescence was carried out using antibodies against the FLAG tag, which is contained in GB1, and COX IV which is a known mitochondrial marker. Nuclei were stained with DAPI. Scale bar: 10 µm.

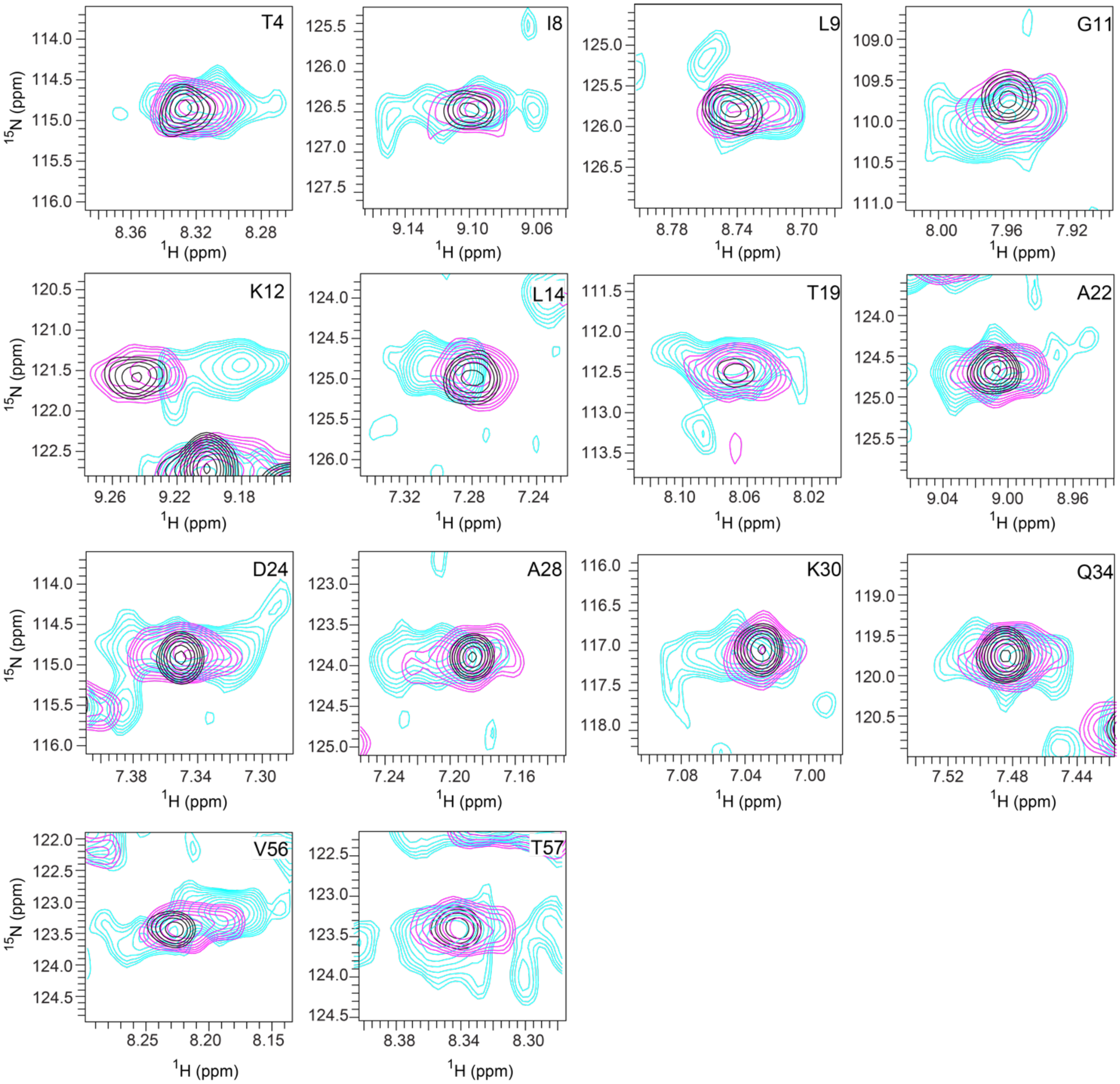


**Fig S9. Mitochondrial localization of mitochondrial-tagged GB1.** GB1 with a mitochondrial tag was purified and electroporated to HEK-293 cells. Spectra acquired immediately after electroporation, at 0 hour, (cyan) is compared to spectra acquired 4 hours post electroporation (magenta). The in-vitro reference spectrum (black) is closer to the average chemical shift of the in-cell NMR spectrum at 4 h (magenta). All the spectra were recorded at 310 K.

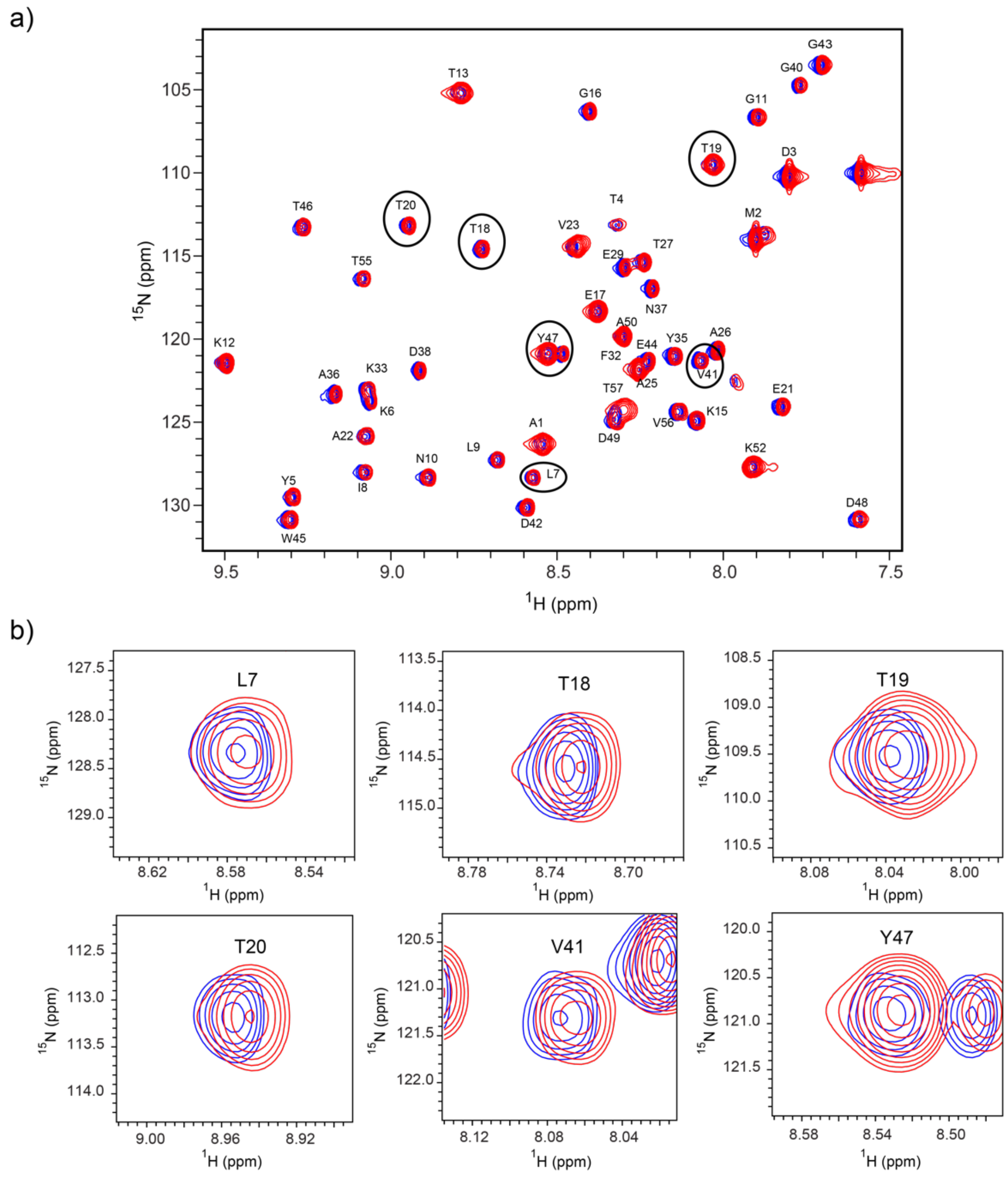


**Fig S10. Effect of purified cell metabolites on the multiplicity of the GB1 resonances.** The peak multiplicity can not be reconstituted by ex situ measurement with GB1 added to cell metabolites. (a) Comparison of GB1 cross peaks acquired in presence of HEK-293 cell metabolites (red) together with the *in vitro* GB1 spectrum (blue). (b) selected resdidues as highlighted in black circles in a) are shown for better visualization of the line shape of cross peaks. For all the residues the cross peaks of the spectra recorded in presence of cell's metabolite do not correlate with the multiplicity of the cross peaks from the in cell NMR spectrum indicating that the peak multiplicity is sensitive to the intact nature of the cell, and interacting to large extent with the cellular proteins and organelles.

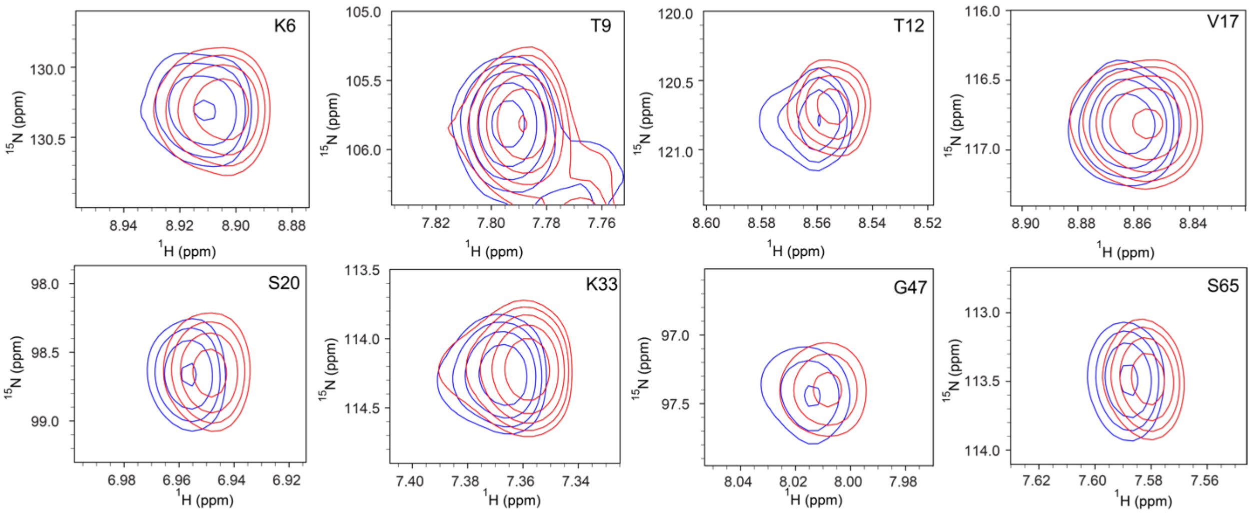


**Fig S11. Effect of purified cell metabolites on the multiplicity of ubiquitin resonances.** The peak multiplicity can not be reconstituted by ex-situ measurement with ubiquitin added to cell metabolites. Selected resonances of ubiquitin spectra acquired in presence of the soluble fraction of the HEK-293 cell metabolites (red) together with the *in vitro* ubiquitin spectrum (blue) are shown.

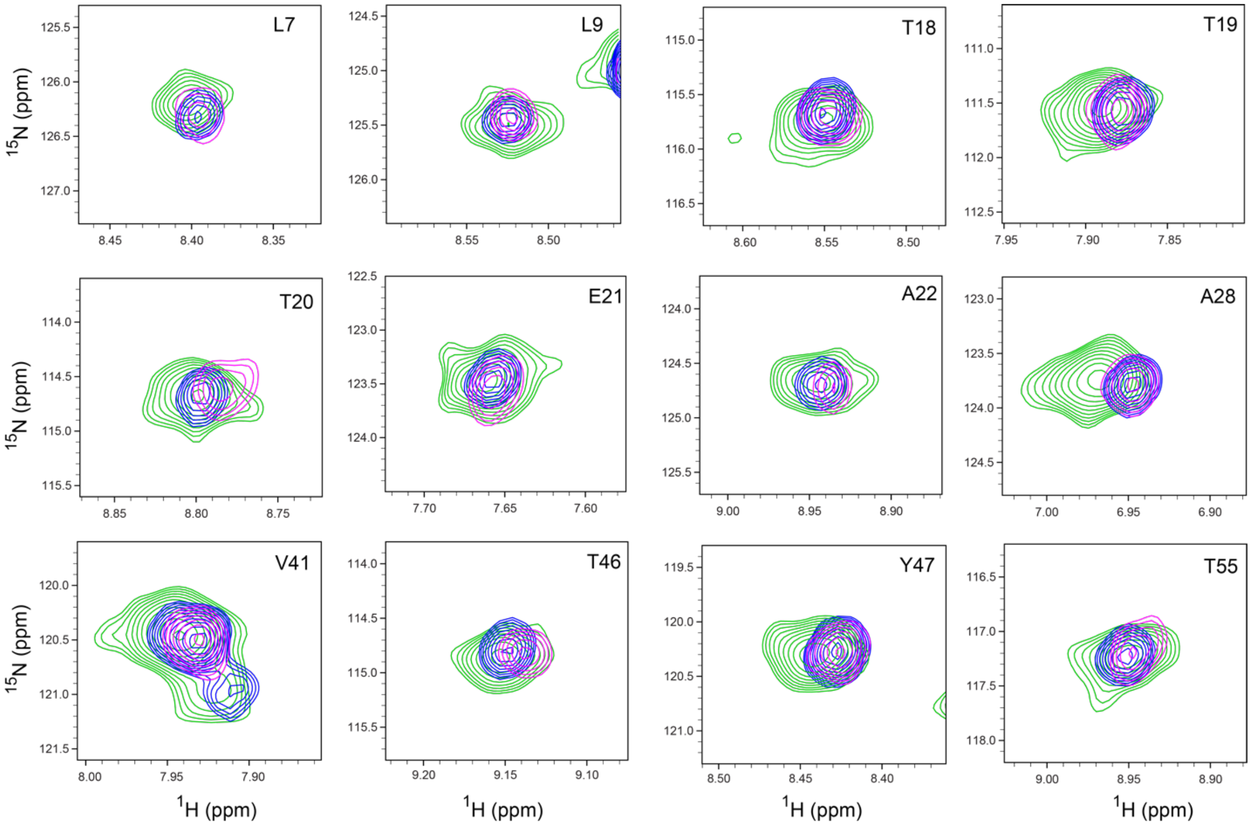


**Fig S12. Concentration dependence of the multiplicity of GB1 cross peaks in cells at 283 K.** Comparison of GB1 cross peaks acquired in living HEK-293 cells with a lower concentration of GB1 (~10 μM, green), with a higher concentration of GB1 (~40 μM, magenta) together with the reference in-vitro spectrum (blue). The contour levels of magenta was adjusted to match that of reference blue spectrum. The rather well superposition between the *in vitro* reference peaks (blue) with the magenta peaks of the spectrum at high GB1 concentration indicate that a saturation occured.

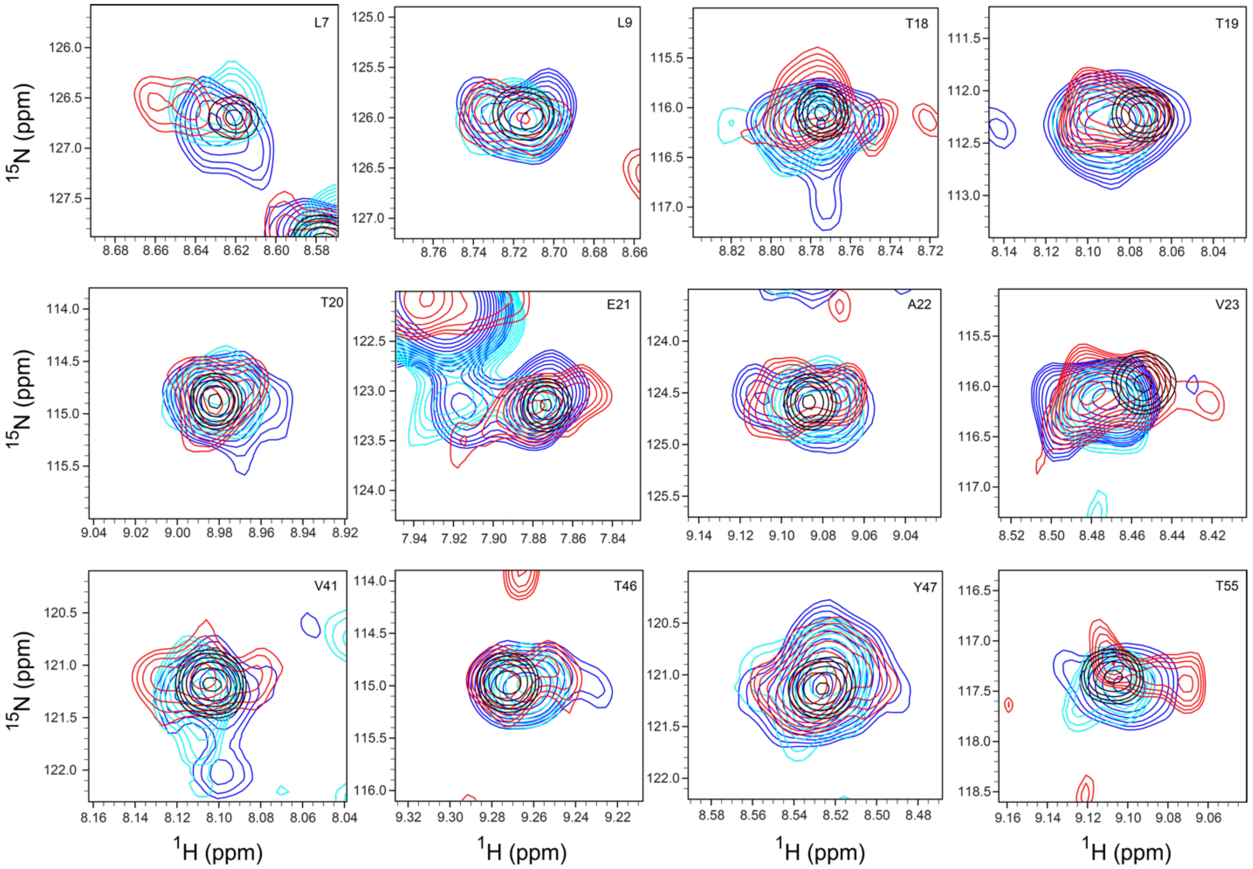


**Fig S13. Concentration dependence of the multiplicity of GB1 cross peaks in cells at 310 K.** Comparison of GB1 cross peaks acquired in living HEK-293 cells with a higher concentration of GB1 (~40 μM, blue), with a lower concentration of GB1 (~10 μM, cyan) together with the reference in-vitro spectrum (black) and in-cell NMR spectrum acquired with ~4 μM of GB1 (red, see Fig. 1). All the spectra were recorded at 310 K.

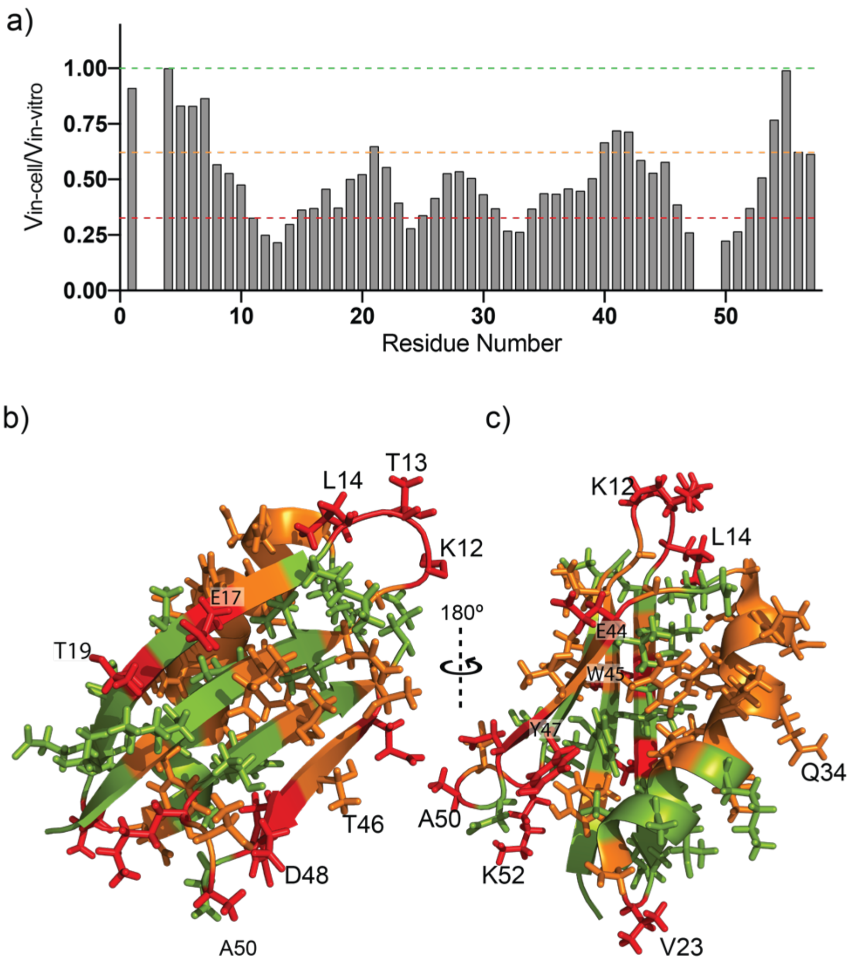


**Fig S14. In-cell NMR revelas peak attenuations on the GB1 resonances.** a) The ratio of the cross peak volume of the NMR resonances from the in cell spectrum to that of the in vitro spectrum are plot versus the sequence. b) and c) Transient interactions identified by signal loss (see a) are plot onto the 3D structure of GB1. Residues are colored red (strong transient interaction), orange (intermediate transient interaction), and green (little transient interaction) based on the extent of reduction in NMR cross peak volume ratio as indicated in a. In particular the dynamics of the loops (e.g. Lys12-Leu14 and Asp48-Lys52) and solvent exposed side chains (e.g. Glu17, Val23, Glu44) are highly affected by the cellular environment while the hydrophobic core (with the exception of the partially solvent exposed Trp45, Tyr35 and Tyr47) seems not to be altered.

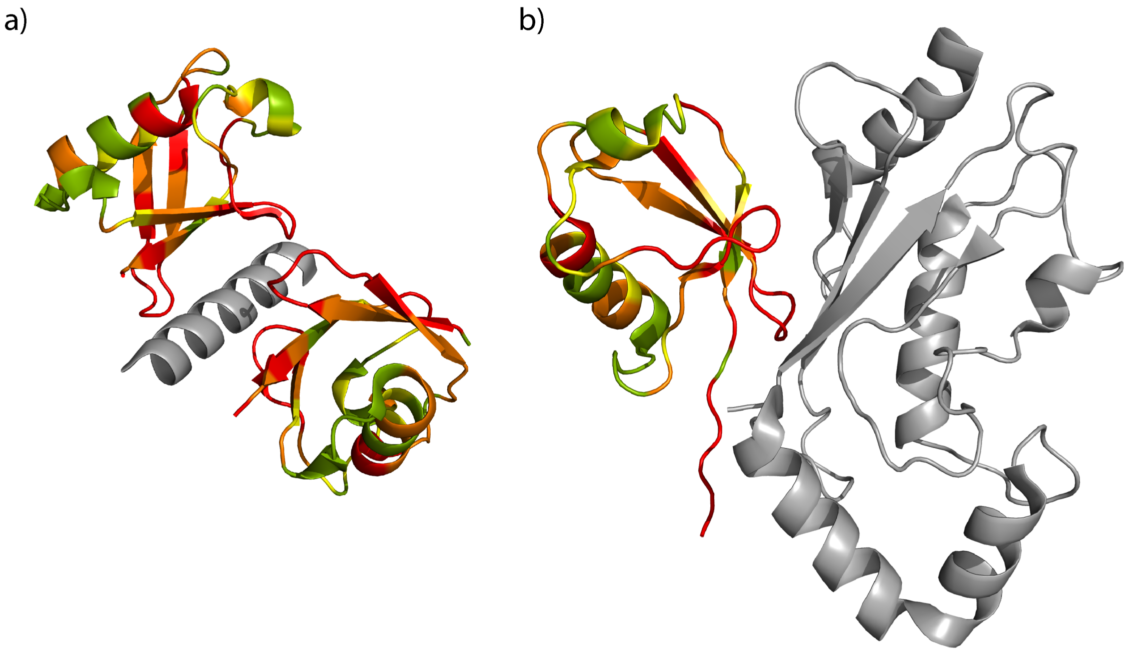


**Fig S15. Structures of ubiquitin–ubiquitin-binding domain (UBD) complexes**. a) PDB 2D3G and b) PDB 2FUH. UBDs are shown in grey. Ubiquitin is shown in the same color code as in the Fig. 2d and 2e to highlight the transient interactions of unknown nature. Peak splitting as well as chemical shifts perturbation and intensity changes of the resonances are observed in cells.

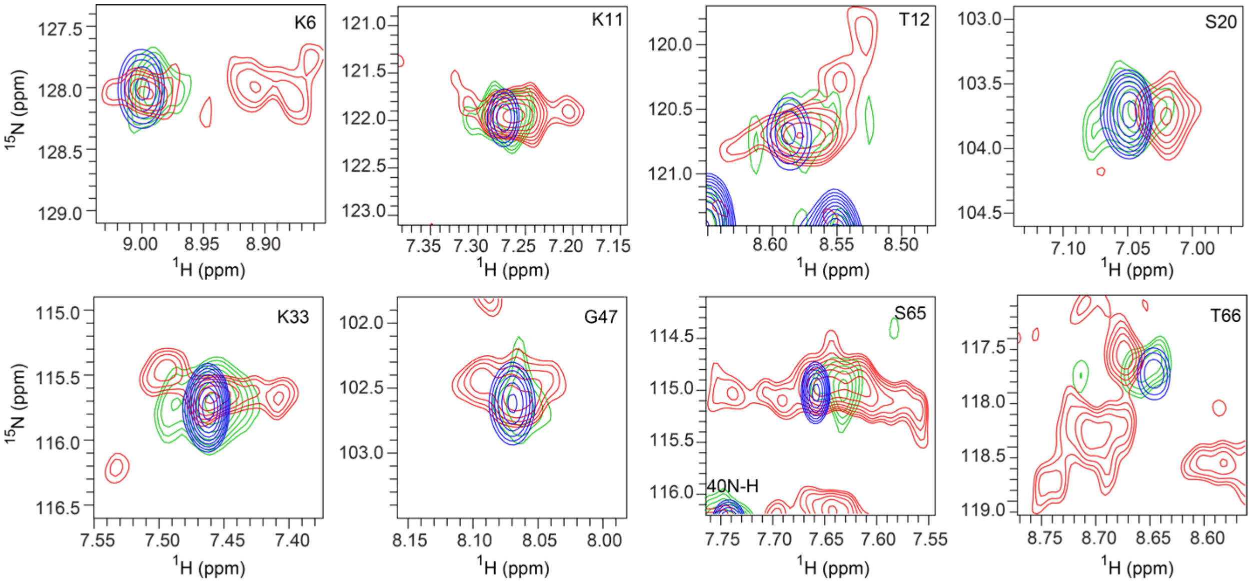


**Fig S16. Inactive ubiquitin obtained by C-terminal deltaGG mutation shows less peak multiplicity in cells when compared with wildtype ubiquitin.** Ubiquitin with a C-terminal deltaGG mutation considered to be an inactive ubiquitin was electroporated into HEK-293 cells and in-cell NMR spectra were acquired as in the case of wild type ubiquitin. Comparison of the peak multiplicity of wild type ubiquitin (shown in red) with its deltaGG variant (green) shows significant reduction in multiplicity. Selected ubiquitin resonances are shown together with the corresponding in vitro ubiquitin resonances (blue).

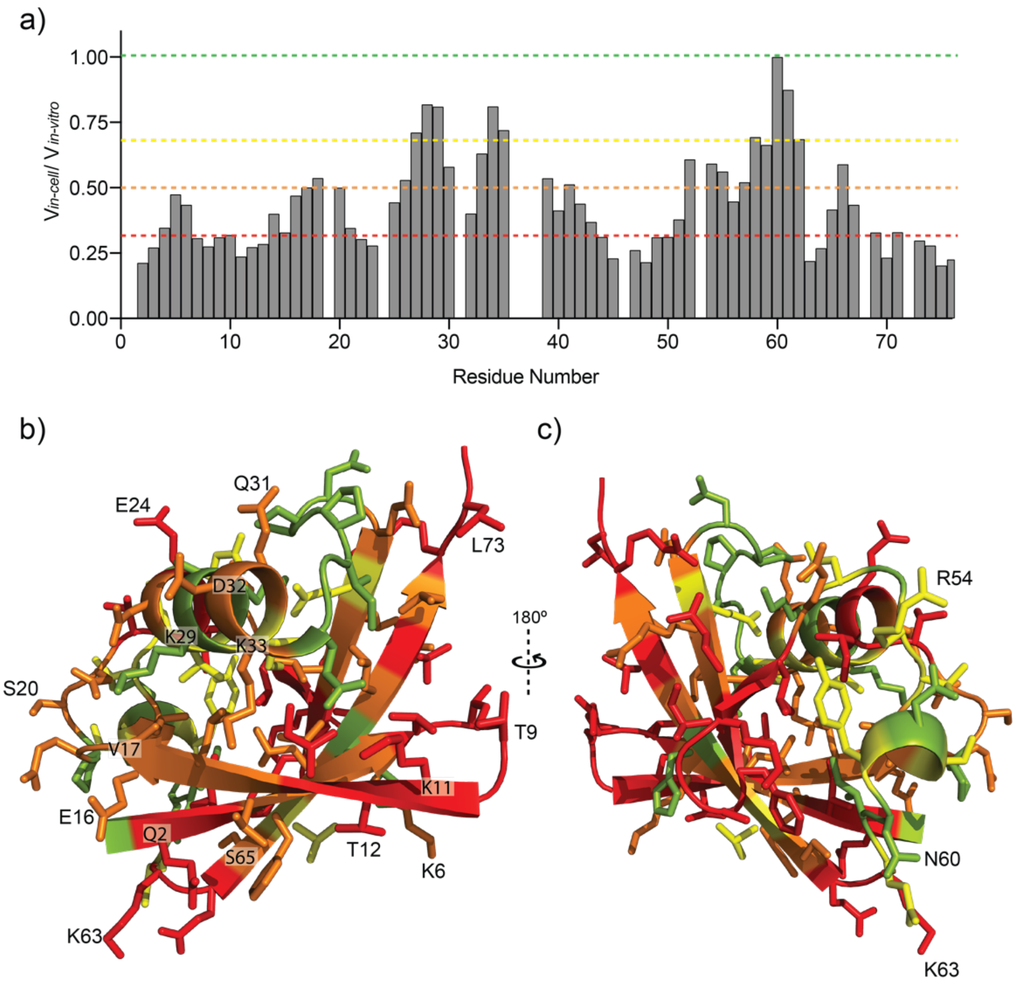


**Fig S17. In-cell NMR revelas peak attenuations on the ubiquitin resonances**.

a) The ratio of the cross peak volume of the NMR resonances from the in cell spectrum to that of the in vitro spectrum are plot versus the sequence. b) and c) Transient interactions identified by signal loss (see c) are plot onto the 3D structure of ubiquitin. Residues are colored red (strong transient interaction), orange (intermediate transient interaction), yellow (slight transient interaction) and green (little transient interaction) based on the extent of reduction in NMR cross peak volume ratio as indicated in a.

Note that all the five $\beta$ strands of ubiquitin which are alternatively anti-parallel and constitute more than half of the 3D structure are significantly affected. In particular the loops 1, 2 and 4 with residue stretches Thr7-Gly10, Val17-Ile23, Phe45-Lys48 undergoes intermediate exchange as evidenced by the reduction in their volume ratios (Fig. S9c). Surprisingly the residues showing transient interactions or/and dynamics induced by the cellular milieu have mainly solvent exposed side chains. In addition to the loops 1, 2 and 4 certain residues in loop 6 such as Lys63 and Glu64 are highly affected by the cellular environment suggesting the involvement of Lys63 in the ubiquitination and transient interactions.

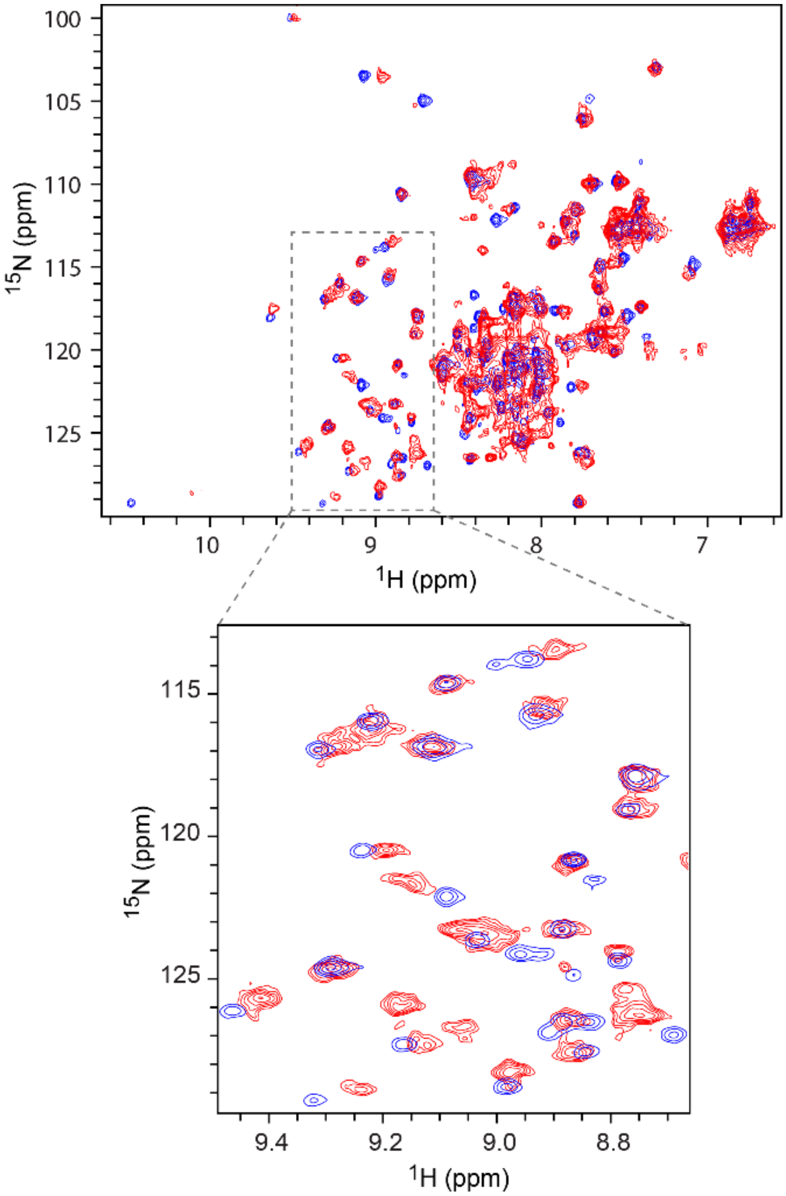


**Fig S18. In cell NMR [$^{15}$N,$^{1}$H]-HMQC spectrum of PDZ2**. Comparison of the in-cell (red) and in-vitro (blue) NMR spectra of PDZ2 domain.

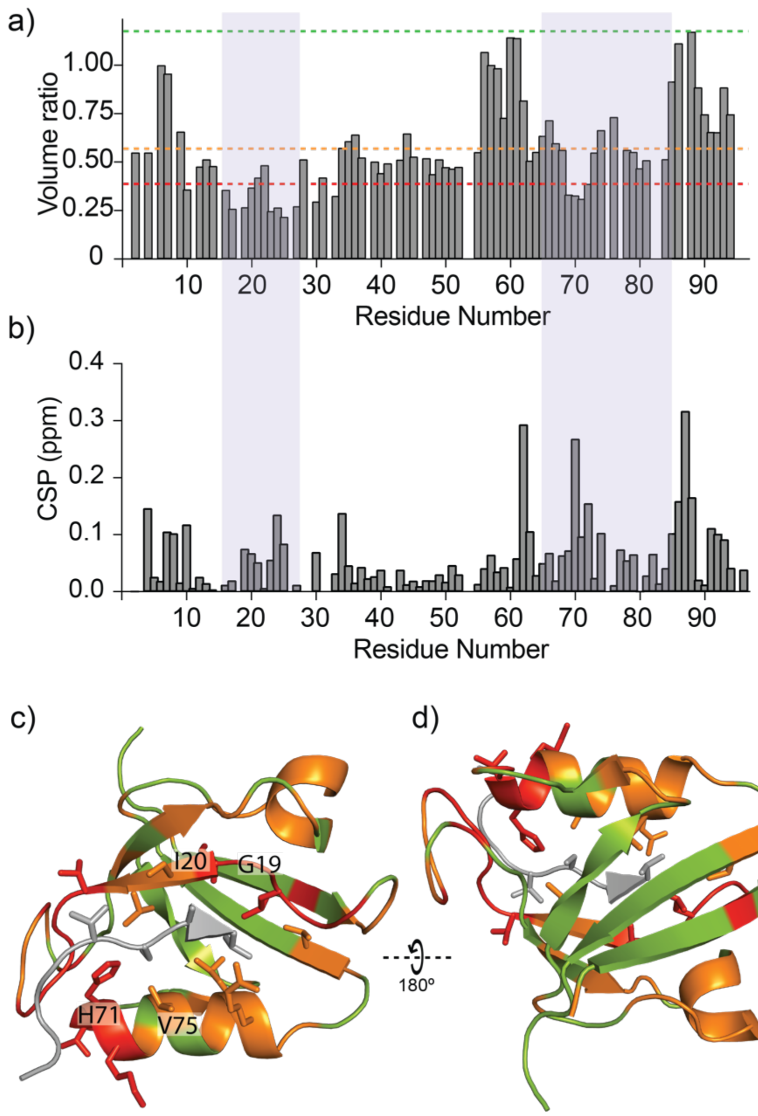


**Fig S19. Transient interaction of PDZ in cells measured by peak volume ratios and chemical shift perturbation**. a) Volume ratio of the in-cell NMR cross peaks of the PDZ2 domain in respect to the in vitro NMR spectrum. A box integration method was used to include multiple peaks of the same residue. b) The chemical shift perturbation of in-cell NMR resonances of PDZ2 domain with respect to the reference in-vitro spectrum. The most affected residues in both plots are highlighted in light blue color. c and d) 3D structure of the PDZ2 domain in complex with a peptide ligand (PDB: 1D5G), where the the ligand peptide colored in grey. Transient interactions identified according to the volume ratio as in Fig. S12a are plot onto the 3D structure of PDZ2 domain. Residues are colored red (strong transient interaction), orange (intermediate transient interaction), and green (little transient interaction) based on the extent of reduction in NMR cross peak volume ratio as indicated in a. Selected residues are labeled.

Note that signal intensity attenuations of the cross peaks of the in-cell NMR spectrum in respect to the in vitro spectrum attributed to transient interactions are observed for the residues at the canonical binding pocket, the short helix comprising residues 47-50 as well as the N/C-terminal $\beta$-strands that are close in space with each other (Fig. 3c,d, Fig S11 and Fig S12c, d).

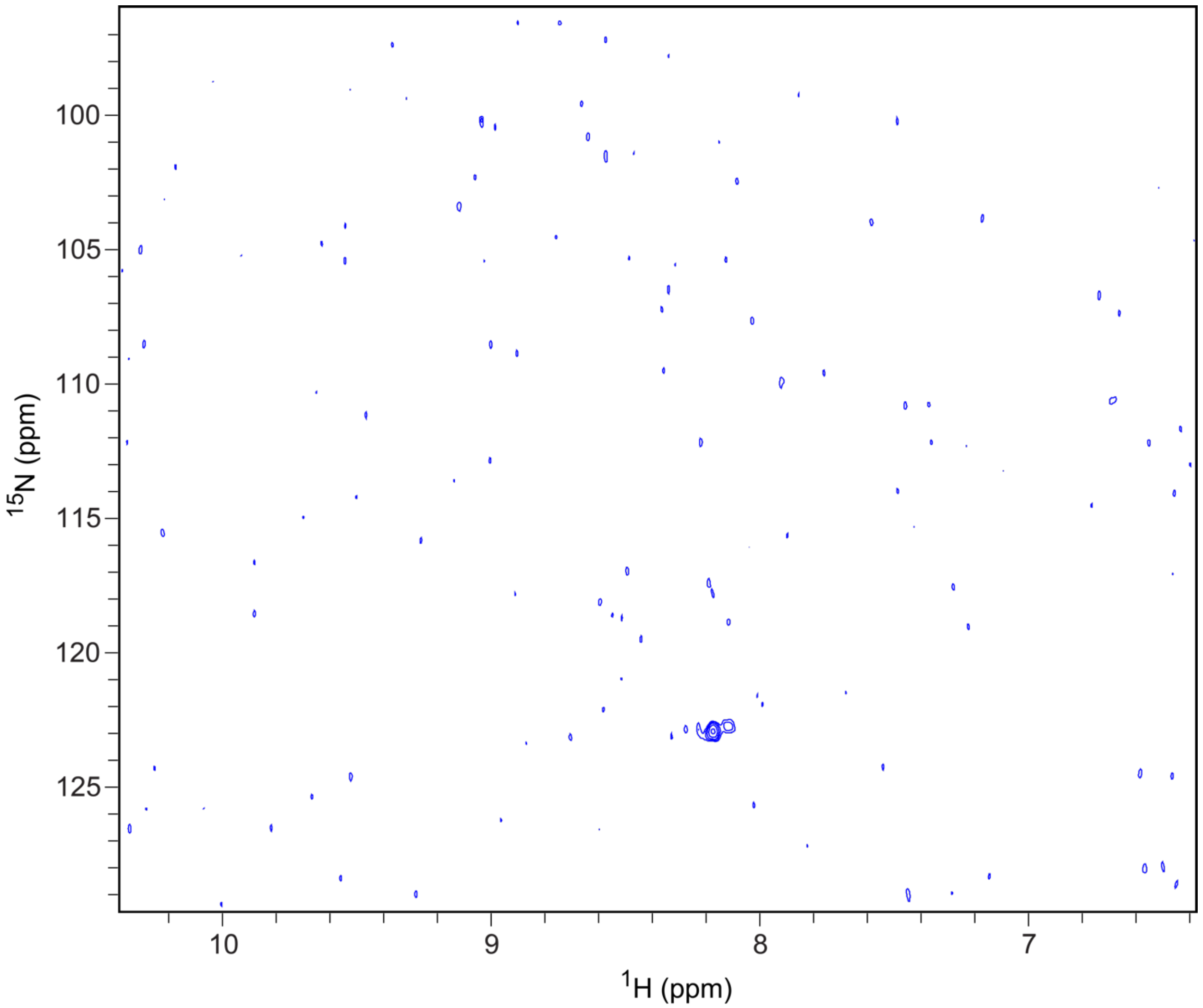


**Fig S20. In-cell NMR spectrum of COS7 cells after empty electroporation.** Standard electroporation procedure was carried out in the absence of isotopically labeled proteins followed by a SOFAST-[$^{15}N$,$^{1}H$]-HMQC NMR experiment as used by the other in cell NMR experimemts. Despite two cross peaks the spectrum is empty indicating a lack of resonances from $^{15}N$-labeled cellular metabolites and other native biomolecules.

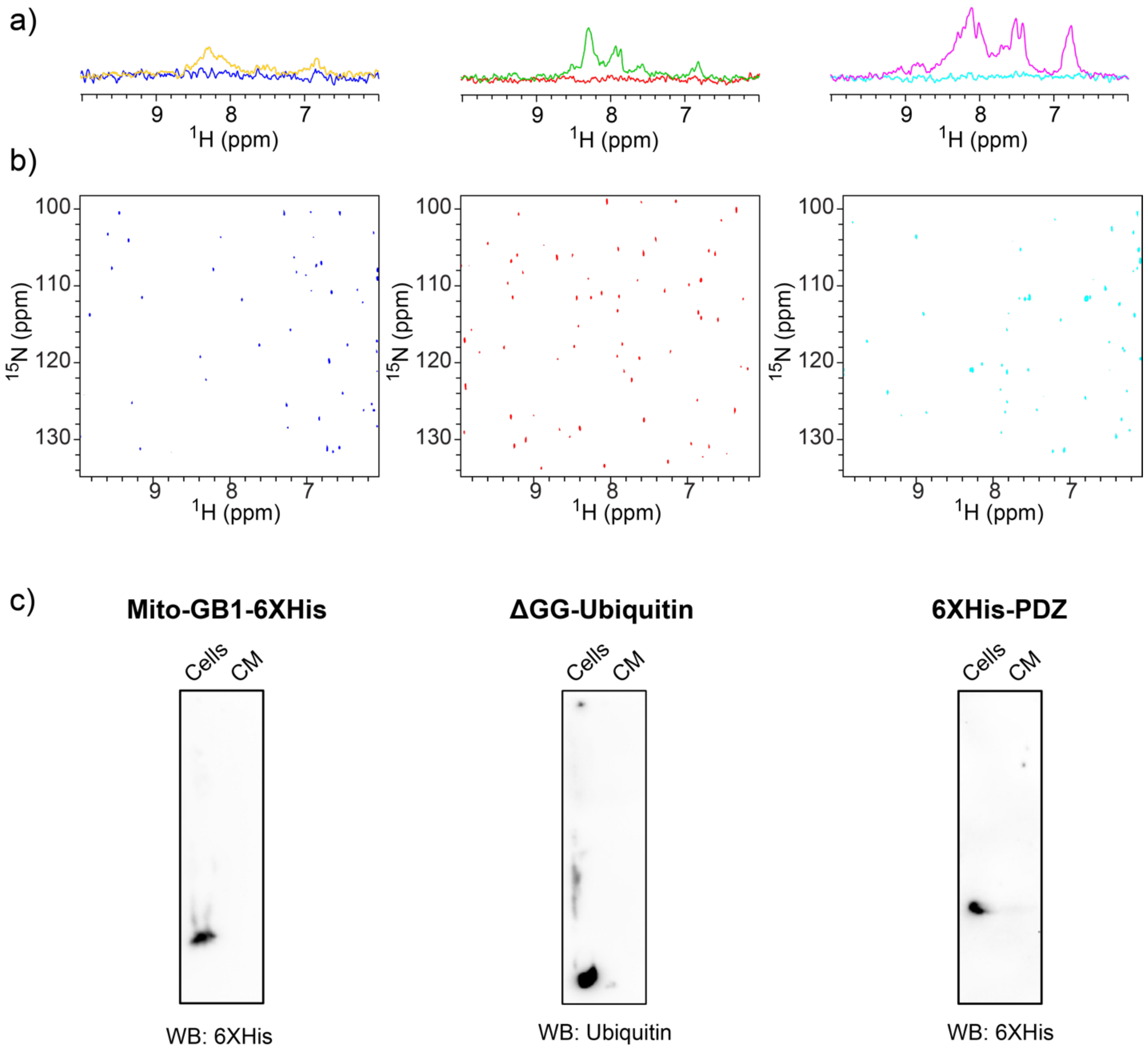


**Fig S21. NMR spectra acquired with in-cell NMR sample and its supernatant after 4h and intact protein identification by Western Blot.** a) Overlay of the $^{15}$N-edited 1D spectra derived from the 2D in-cell NMR spectra and spectra of the corresponding supernatant of GB1 (left), Ubiquitin (middle) and PDZ2 domain (right) 4 h post electroporation. 1D spectra with orange, green and magenta colors correspond to the in-cell NMR spectra and those with blue, red and cyan correspond to the cell supernatant. (b) The 2D spectra of the cell supernatant 4 h post electroporation with color coding with respect to Fig. S21a corresponds to GB1, Ubiquitin and PDZ2 domain, respectively. (c) The western blot images of GB1, ubiquitin, and PDZ domain using the indicated antibodies show no sign of protein degradation.

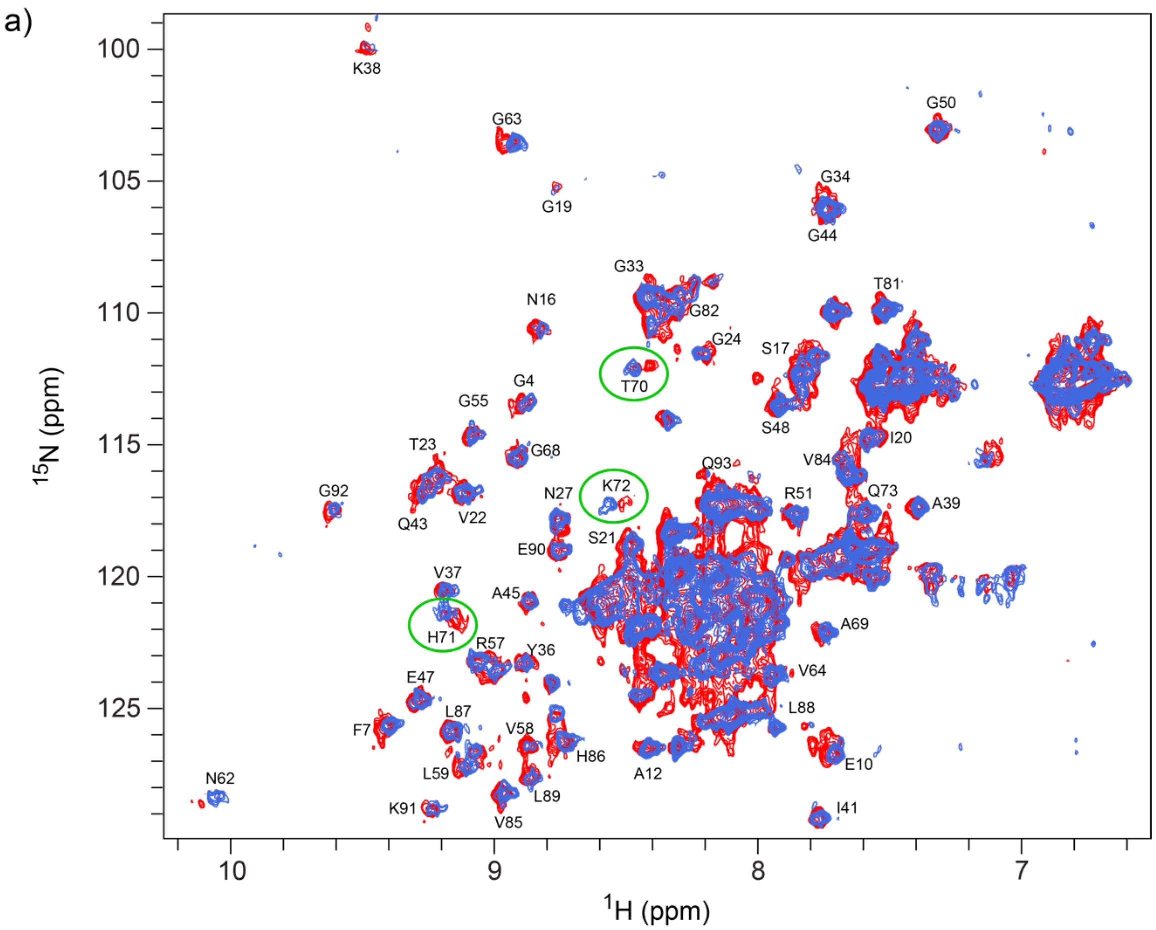


**Fig S22. In-cell NMR spectra of PDZ domain in COS7 cells at 4 and 8 hours.** The spectra collected 4 h (blue) and 8h (red) post electroporation are comparable except for the residues next to the pH sensitive His71 such as T70, H71 and K72 highlighted by green circles. They show pH dependent chemical shift perturbations attributed to the decrease of the pH caused by cell metabolism.
